# *Pantoea ananatis* defeats *Allium* chemical defenses with a plasmid-borne virulence gene cluster

**DOI:** 10.1101/2020.02.12.945675

**Authors:** Shaun P. Stice, Kyle K. Thao, Chang Hyun Khang, David A. Baltrus, Bhabesh Dutta, Brian H. Kvitko

## Abstract

Onion (*Allium. cepa* L), garlic (*A. sativum* L.), and other members of the *Allium* genus produce volatile antimicrobial thiosulfinates upon cellular damage. Allicin has been known since the 1950s as the primary antimicrobial thiosulfinate compound and odorant produced by garlic. However, the roles of endogenous thiosulfinate production in host-bacterial pathogen interactions have not been described. The bacterial onion pathogen *Pantoea ananatis*, which lacks both the virulence Type III and Type II Secretion Systems, induces necrotic symptoms and extensive cell death in onion tissues dependent on a proposed secondary metabolite synthesis chromosomal gene cluster. We found strong correlation between the genetic requirements for *P. ananatis* to colonize necrotized onion tissue and its capacity for tolerance to the thiosulfinate allicin based on the presence of an eleven gene, plasmid-borne, virulence cluster of sulfur/redox genes. We have designated them ‘*alt*’ genes for <u>al</u>licin tolerance. We show that allicin and onion thiosulfinates restrict bacterial growth with similar kinetics. The *alt* gene cluster is sufficient to confer allicin tolerance and protects the glutathione pool during allicin treatment. Independent *alt* genes make partial phenotypic contributions indicating that they function as a collective cohort to manage thiol stress. Our work implicates endogenous onion thiosulfinates produced during cellular damage as mediators of interactions with bacteria. The *P. ananatis*-onion pathosystem can be modeled as a chemical arms race of pathogen attack, host chemical counter-attack, and pathogen resistance.

**Significance Statement:** Alliums (e.g. onion and garlic), after sustaining cellular damage, produce potent antimicrobial thiosulfinates that react with cellular thiols. The bacterial onion pathogen *Pantoea ananatis*, which lacks the virulence Type III and Type II Secretion Systems, induces cell death and necrotic symptoms on onions. We have identified a plasmid-borne cluster of sulfur/redox virulence genes that 1) are required for *P. ananatis* to colonize necrotized onion tissue, 2) are sufficient for tolerance to the thiosulfinates, and, 3) protect the glutathione pool during thiosulfinate treatment. We propose that the thiosulfinate production potential of *Allium* spp. governs *Allium*-bacterial interaction outcomes and that the *P. ananatis*-onion pathosystem can be modeled as a chemical arms race of attack and counterattack between the pathogen and host.

## Introduction

Plants deploy diverse chemical weaponry to defend themselves from microbial pathogens and herbivorous pests. Among these forms of chemical defenses, there are several examples of potent reactive antimicrobials and toxins that are converted into their active forms only after plant tissue damage. Cellular damage allows preformed precursor substrates and activating enzymes stored in separate sub-cellular compartments to mix and react (1). These damage-driven enzymatic and chemical reactions allow rapid, potent, and often volatile, reactive chemical responses to tissue damage associated with herbivory and disease necrotrophy. Charismatic examples include glucosinolates that give brassicaceaous vegetables their distinct pungency as well as the cyanogenic glycosides that accumulate in rosaceous seeds (2, 3). Onion (*Allium cepa* L.), garlic (*Allium sativum* L.), and other alliaceous spp. display a classic example of this form of defense producing an array of reactive organosulfur compounds through a combination of enzymatic and chemical reactions upon tissue damage (1, 4). These compounds are associated with the characteristic flavors and odors of onion and garlic and also act as volatile irritants and antimicrobial compounds (5, 6). The thiosulfinate allicin (diallyl thiosulfinate, structure 3a), produced by the action of alliinase (EC 4.4.1.4) on alliin (*S*-2-propenyl-L-cysteine sulfoxide, structure 1a), is responsible for the odor of crushed garlic and is the primary volatile antimicrobial compound in garlic extract (7, 8). Allicin acts as a thiol toxin, reacting with cellular thiols and producing allyl-mercapto protein modifications, which can directly inactivate enzymes and cause protein aggregation (9). Allicin also reacts with reduced glutathione converting it into allylmercaptoglutathione and thereby depleting the reduced glutathione pool (7, 9). Asymmetric 1-propenyl methyl thiosulfinates (structure 3b) accumulate after disruption of onion tissue with total thiosulfinate production in onion extracts in the range of 100s of nmol per gram fresh weight (10). In onion, the combined action of alliinase and a second enzyme, lachrymatory factor synthase (LFS) converts the majority of isoalliin (*S*-1-propenyl-L-cysteine sulfoxide, structure 1c) into syn-propanethial-S-oxide (a.k.a. lachrymatory factor, structure 4), the chemical irritant of onion associated with inducing tears (11). Conversely, garlic, which lacks LFS, accumulates nearly 100-fold higher total thiosulfinate per gram fresh weight than onion (10).

Onion center rot, caused by at least four species bacteria in the *Pantoea* genus from the order Enterobacterales, is an economically impactful disease of onions that routinely results in significant losses to yield and marketability (12, 13). In the southeastern United States, onion center rot is primarily caused by *Pantoea ananatis*. No onion cultivars with resistance to pathogenic *P. ananatis* have yet been identified*. P. ananatis* invades leaves through wounds and causes blights and wilting of infected leaves. The pathogen can also invade bulbs through infected foliar tissues. Bulb invasion is associated with systemic movement of the pathogen from the leaf blade to the corresponding scale in the onion bulb (14). Preferential feeding by multiple species of thrips insects (*Frankliniella fusca*, and *Thrips tabaci)* around the onion neck, has been observed to play a role in transmission and dissemination of this pathogen from epiphytic onion or weed populations into onion foliage (15, 16).

*Pantoea ananatis* is a broad-host-range pathogen able to cause disease in diverse monocot hosts including onion, pineapple, maize, rice, and sudangrass (17). *P. ananatis* is also a rare example of a gram negative plant pathogen that lacks both a virulence-associated Type III Secretion System (T3SS) used by many plant pathogenic bacteria to deliver immune-dampening effector proteins as well as a Type II Secretion System (T2SS) commonly used to deliver plant cell wall degrading enzymes associated with soft rot pathogens (18, 19). This surprising lack of the key pathogenicity factors, typically associated with gram negative bacterial plant pathogens, has left the primary molecular mechanisms by which *P. ananatis* causes disease an open question.

Recent work by Asselin *et al.* and Takikawa *et al*. identified the horizontally transferred chromosomal gene cluster HiVir (<u>Hi</u>gh <u>Vir</u>ulence, also known as PASVIL, *<u>P</u>antoea <u>a</u>nanatis* <u>s</u>pecific <u>vi</u>rulence <u>l</u>ocus) as a critical pathogenicity factor for *P. ananatis* to produce necrotic symptoms on onion (20, 21). HiVir is hypothesized to encode a biosynthetic gene cluster for an, as of yet, undescribed secondary metabolite that may act as a plant toxin. The metabolite synthesized by the HiVir cluster is predicted to be a phosphonate or phosphinate compound based on the presence of a characteristic *pepM*, phosphoenolpyruvate mutase gene in the HiVir cluster that is essential for necrosis induction (20).

Upon induction of HiVir-dependent necrosis, it is expected that onion tissues would become a noxious and challenging environment for microbial colonization due to the endogenous production of reactive sulfur antimicrobial compounds. In prior comparative genomics analysis of *P. ananatis*, we identified four clusters of plasmid borne genes, OVRA-D, that correlated with onion virulence (19). Using mutational analysis, we identified an eleven gene sub-cluster within OVRA that is critical for colonization of onions during disease as well as growth in necrotized onion bulb tissue and onion extracts. This cluster is comprised of genes with annotations associated with sulfur metabolism and redox and confers tolerance to the thiosulfinate allicin. We determined that onion extract contains thiosulfinates at concentrations sufficient to restrict bacterial growth and restricts *Pantoea* growth with similar kinetics to allicin. We designated these genes *alt* for <u>al</u>licin tolerance. Expression of the *alt* cluster conferred virulence capacity and allicin tolerance to a natural *P. ananatis* isolate lacking *alt* genes and also conferred allicin tolerance to *E. coli*. We observed that the *alt* genes likely encode an additive cohort of protective enzymes to manage cellular thiol stress with multiple genes independently conferring partial phenotypes. These results demonstrate that onion thiosulfinates play an important role in biotic interactions with bacteria and that *P. ananatis* pathogenic interactions with onion can be modeled as a chemical arms race based on pathogen attack, host counterattack, and specialized pathogen adaptations for chemical defense.

## Results

### Plasmid-borne OVRA genes promote onion scale colonization and facilitate growth in onion extract

Onion pathogenic *P. ananatis* causes tissue necrosis on onions in a cultivar-independent manner (19, 20, 22, 23). Recently, Asselin *et al*., identified the proposed biosynthetic gene cluster HiVir carrying a *pepM* gene essential for the production of both onion leaf and bulb necrosis (20). We previously developed an assay for screening *P. ananatis* pathogenesic potential based on the development of a clearing zone around the inoculation site of red onion scales (19). The red scale clearing phenotype required the HiVir *pepM* gene similar to other necrosis phenotypes (Fig 1A, 1B). We determined that the clearing zone in red onion scales is associated with extensive onion cell death although plant cell walls appear to be left intact. This observation was based on apparent plasmolysis and the staining of nuclei by propidium iodide, indicating loss of plasma membrane integrity, which was consistent with the lack of staining with the vital stain fluorescein diacetate (24, 25). This is in stark contrast to observations of cells from non-cleared tissue in the same scale samples (Fig. 1C).

**Fig 1.**
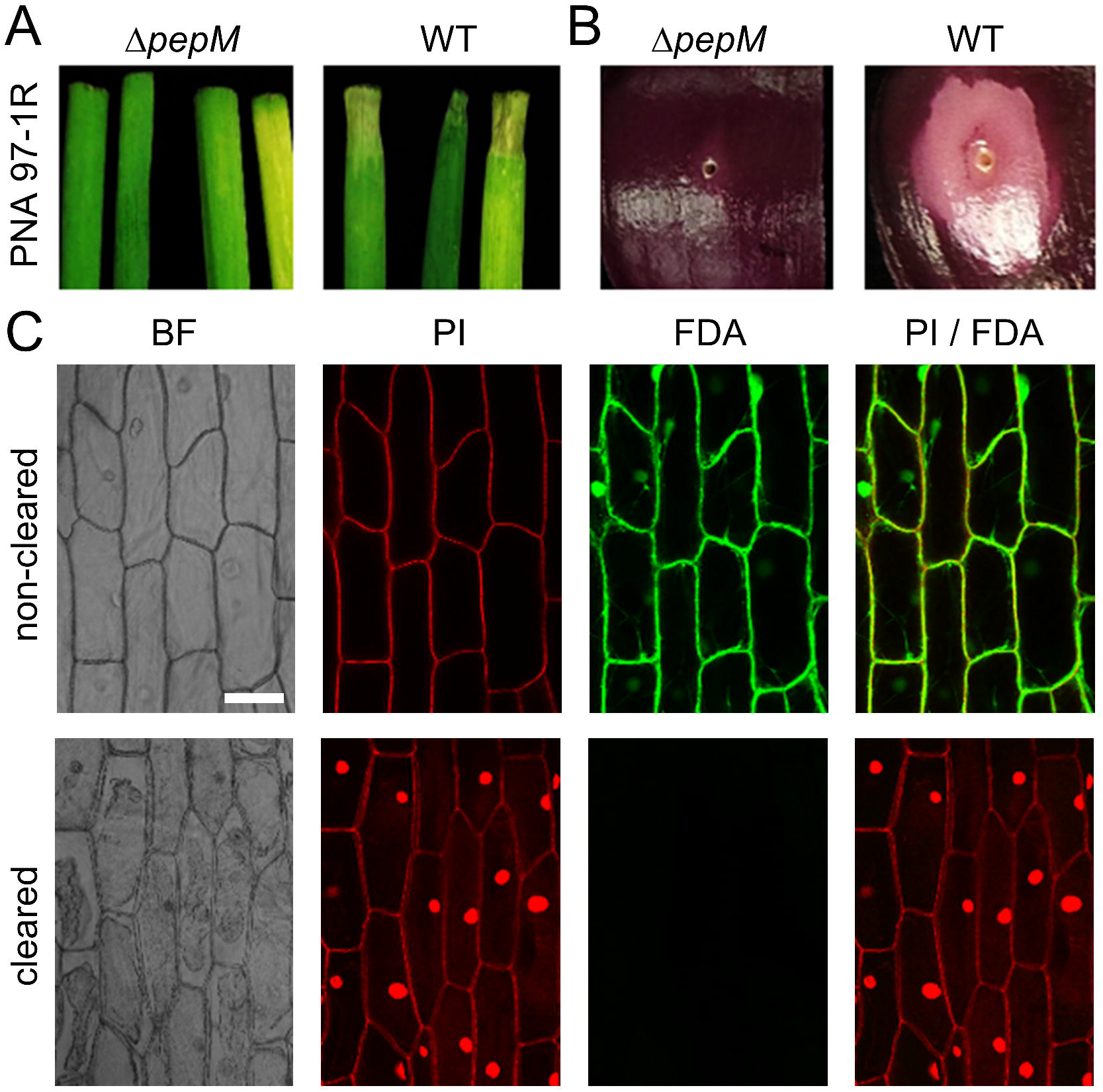
The HiVir *pepM* gene is required for inducing cell death in red onion scales. (*A*) ∆*pepM* mutant in PNA 97-1R background does not cause foliar lesions on onion blades. Images were taken 3 DPI on 6-week-old onion seedlings (cv. ‘century’). Images are not to scale. (*B*) ∆*pepM* does not induce red onion scale clearing. Images were taken 3 DPI and are not to scale. (C) Confocal images of onion epidermal cells in non-cleared and cleared regions stained to determine cellular integrity. BF(grayscale), bright field; PI (red), propidium iodide; FDA(green), fluorescein diacetate. Bar = 100 µm.

In our previous comparative genomics analysis we identified 57 genes in four contiguous blocks on a 161-KB megaplasmid that were strongly correlated with *P. ananatis* virulence on onion (NBCI accession CP020945.2) (Fig. 2A) (19). We sought to determine whether these plasmid-borne OVR (<u>O</u>nion <u>V</u>irulence <u>R</u>egions) genes contributed to *P. ananatis-*mediated onion center rot. Using allelic exchange, we generated deletion mutants of the OVRA, OVRB, OVRC, and OVRD clusters in *P. ananatis* PNA 97-1R (WT) (*SI Appendix, Materials and Methods*). We observed that the OVRA cluster deletion strain both produced smaller clearing zones in a red scale necrosis assay (Fig. 2B and 2C) and reached two log-fold lower bacterial load in onion scale tissue compared to the WT strain and other OVR deletions (Fig. 2C). As PNA 97-1R WT was able to grow to high loads in the dead onion tissue of scale clearing zones, we posited that bacterial growth in clarified, filter sterilized red onion juice (red <u>o</u>nion <u>e</u>xtract: ROE) would be effective proxy for extreme onion tissue damage and that monitoring growth capacity of *P. ananatis* in ROE would mimic colonization potential in dead onion tissue. We monitored bacterial growth in ROE as the change in OD_600_ over time. PNA 97-1R was unable to grow in full strength ROE. However, in half strength ROE, WT, ∆OVRB, ∆OVRC, and ∆OVRD strains grew well while the ∆OVRA strain had a dramatic growth defect (Fig. 2D). We considered two hypotheses for why a ∆OVRA strain lost the capacity for efficient growth in ROE and necrotized onion tissue 1) ∆OVRA lost the ability to utilize key nutrients from host tissue or, 2) ∆OVRA lost the ability to tolerate onion inhibitory factor(s). To test the nutritional model we assessed bacterial growth in a 1:1 mixture of ROE and LB media. The ∆OVRA strain still displayed a strong growth defect in ROE:LB (Fig. 2D). The reduced growth of a ∆OVRA strain under nutritionally replete conditions indicated that the onion inhibitory factor model was more probable. We observed similar growth patterns in ROE and ROE:LB by natural variant *Pantoea* isolates based on the presence or absence of OVRA genes (Fig. S1).

**Fig 2.**
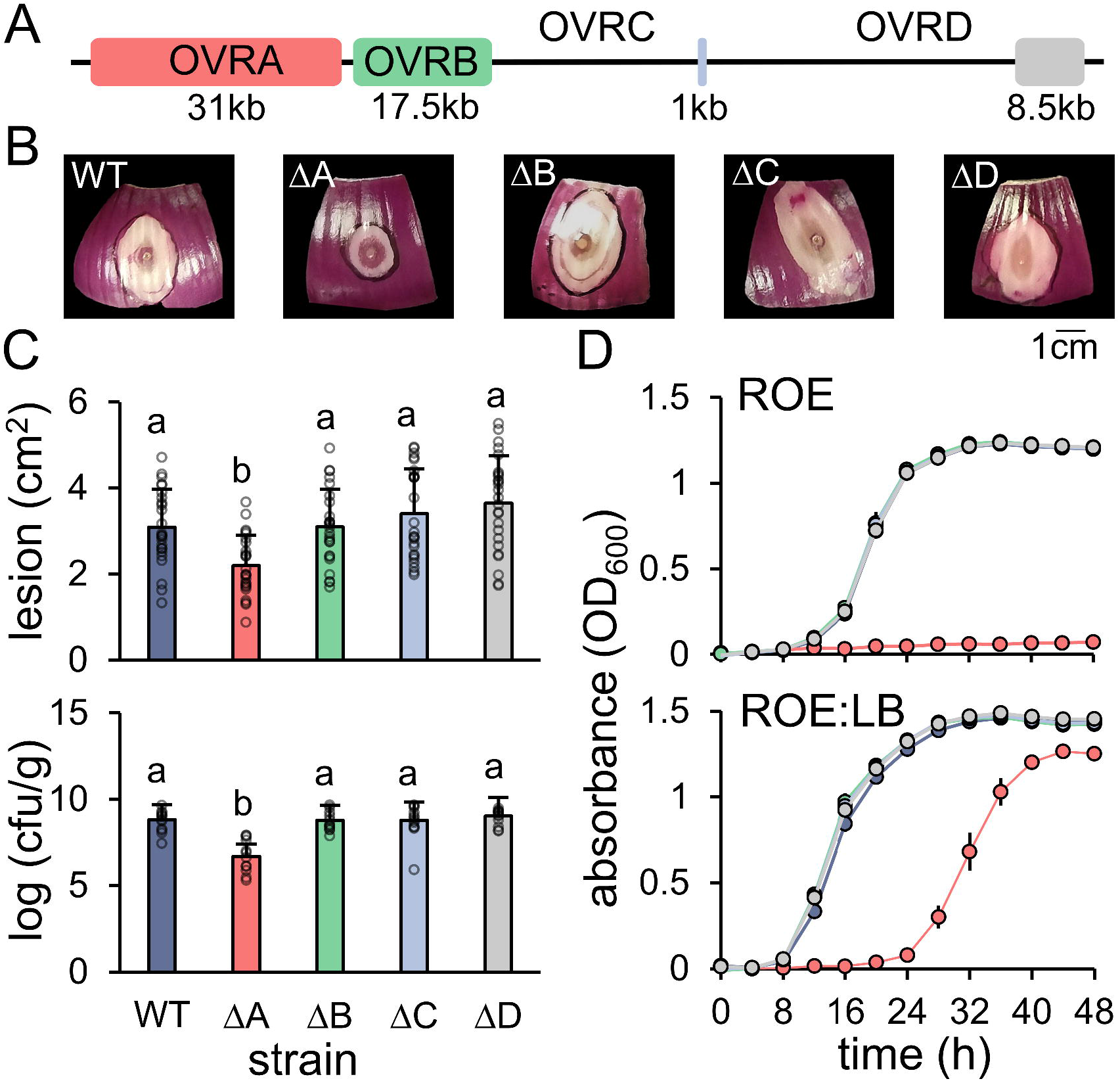
Plasmid-borne OVRA genes promote onion scale colonization and facilitate growth in onion extract. (*A*) Linear graphical representation of the <u>O</u>nion <u>V</u>irulence <u>R</u>egion (OVR) gene clusters on the pOVR mega-plasmid of *P. ananatis* PNA 97-1R (NCBI accession CP020945.2). (*B*) Representative bacterial lesions produced on red onion scales. Image was taken three days post inoculation. (*C*) PNA 97-1R ∆OVRA produces smaller lesions (top, *N=24*) and reaches lower bacterial loads in onion scale tissue (bottom, *N=12*). The data from three independent replicates is presented (one-way ANOVA followed by Tukey’s post-test, *p*<0.001, letters represent significant dif). Error bars represent ±SD. Log_10_ (cfu/g), colony-forming units per gram of onion scale tissue. (*D*) PNA 97-1R ∆OVRA growth (change in OD_600_, Bisocreen C) in liquid culture is inhibited in an aqueous red onion extract (ROE) and delayed when supplemented with LB. This experiment was repeated three times with similar results (*N=6*). Error bars represent ±SEM.

### The 11 gene *alt* sub-cluster in OVRA confers tolerance to allicin, protects the glutathione pool during allicin treatment, and promotes onion bulb colonization

Based on the hypothesis that onion inhibitory factors could be onion thiosulfinates, or possibly another sulfur compound, we focused on the eleven contiguous OVRA genes annotated for functions related to sulfur metabolism and redox (Fig. 3A). Based on various gene annotation pipelines, these 11 genes encode a TetR-family repressor, four reductases including a glutathione disulfide reductase, two potential peroxidases, a thioredoxin and a thioredoxin-like gene, a carbon sulfur lyase and a cysteine-transporter family protein (Table S1). We used allelic exchange to delete the 11 gene sulfur metabolism/redox cluster from OVRA. Using zone of inhibition assays as well as liquid media growth assays, we observed that the OVRA sub-deletion had increased sensitivity to the thiosulfinate allicin (Fig. 3B, Fig S2A) as well as to garlic extract (Fig. S2B). Thus, we chose to name this 11 gene region the *alt* cluster for <u>al</u>licin tolerance. We had noted previously that the sequenced *Enterobacter cloacea* isolate EcWSU1, which is also an onion bulb pathogen, caries cluster of plasmid borne genes similar to the *Pantoea alt* cluster (Fig. S3) (19). Similar patterns of allicin and garlic extract sensitivity were observed in natural variant *Pantoea* isolates based on the presence or absence of OVRA genes or the *alt*-like cluster in *Enterobacter* isolates (Fig. S4). In addition, three chromosomal clusters conferring allicin tolerance via heterologous expression were recently described from the non-pathogenic garlic saprophyte *Pseudomonas fluorescens Pf*AR-1 (26). These clusters from *Pf*AR-1 share similar gene content to the *alt* cluster although not fine scale synteny (26). We observed that a ∆OVRB/C/D mutant showed similar levels of allicin tolerance to WT PNA 97-1R (Fig. 3B). However, we also observed that the ∆OVRA/B/C/D mutant had higher allicin sensitivity than the Δ*alt* mutant. (Fig. 3B). This indicates that some OVR genes outside 11 gene *alt* gene cluster contribute to full allicin tolerance in PNA 97-1R.

**Fig 3.**
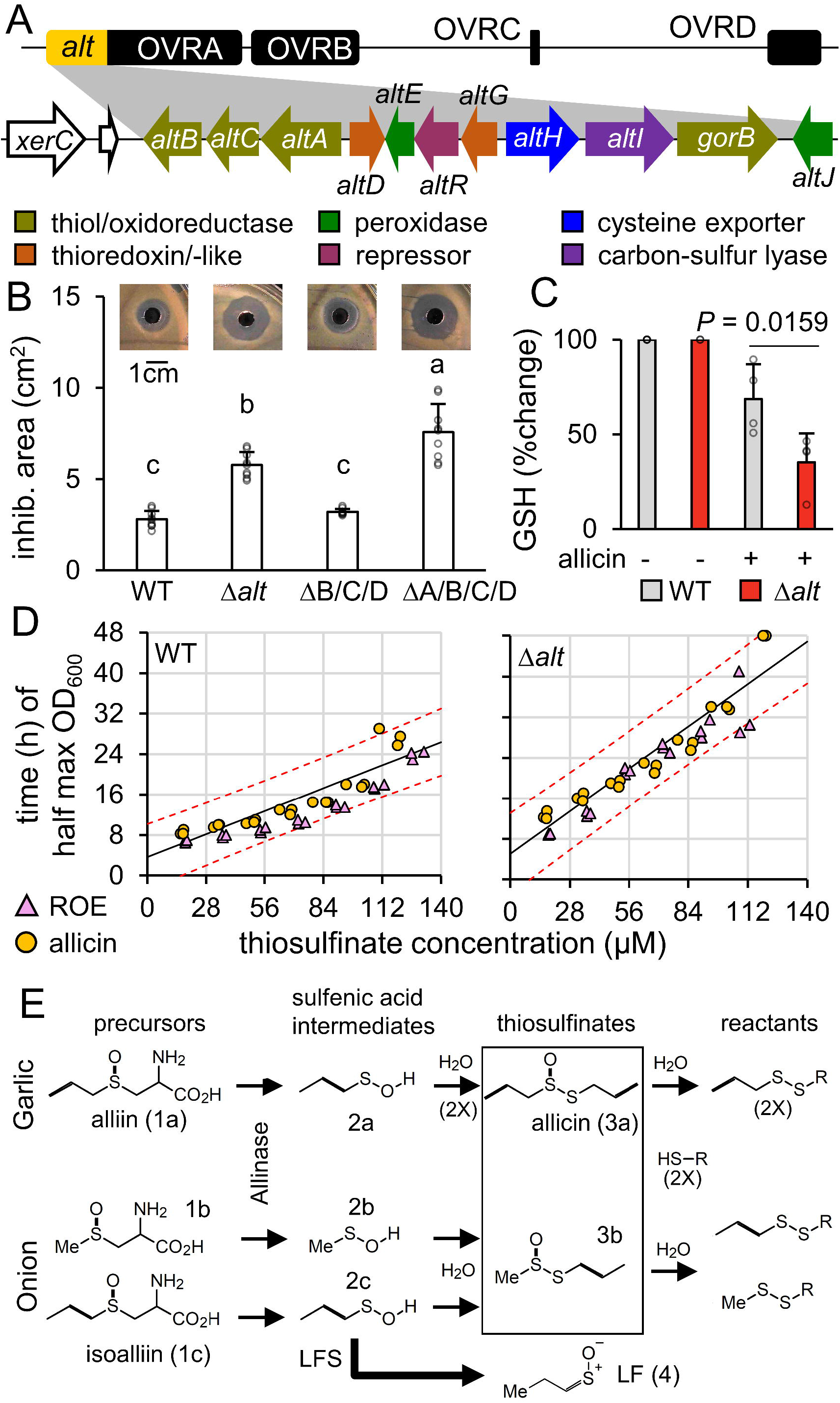
The 11 gene alt sub-cluster in OVRA is critical for tolerance to allicin and onion thiosulfinates. (*A*) Linear graphical representation of the OVR gene clusters of PNA 97-1R mega-plasmid expanded to present 11 genes in the *alt* cluster. Genes are color coded to proposed function. (*B*) Allicin inhibition area of PNA 97-1R mutants. PNA 97-1R ∆*alt* and quadruple mutant PNA 97-1R ∆OVRA/B/C/D have increased susceptibility to allicin compared to PNA 97-1R WT and triple mutant PNA 97-1R ∆OVRB/C/D. This experiment was repeated three times with similar results (*N=6*, one-way ANOVA followed by Tukey’s post-test, *p*<0.001, letters represent significant dif). Error bars represent ±SD. Representative images of clearing zones are included above respective bars. (*C*) The percent change in total glutathione comparing untreated to 1 h allicin stress treatments. PNA 97-1 ∆*alt* has significantly lower levels of glutathione as a percent of its paired untreated sample compared to PNA 97-1R WT. The data from four independent replicates is presented (*N=4*, t-test). Error bars represent ±SE. (*D*) WT and ∆*alt* 48 h thiosulfinate growth response curves. A dilution series of allicin in LB was generated and the thiosulfinate content was measured (4-MP assay) and is presented on the x-axis. The time point at which the culture reached half max OD_600_ (Bioscreen C) during 48 h was is presented on the y-axis (yellow circle). A linear regression of the allicin growth response (solid black line, [WT] y=0.162x+3.64, [∆*alt*] y=0.23x+5.08) was plotted along with 95% confidence intervals (dotted red line). The thiosulfinate concentration and growth response of a ROE dilution series in LB is plotted over the allicin response curve (pink triangle). This experiment was repeated three times. Each point represents an independent bisocreen well. (*E*) Simplified thiosulfinates reaction pathways from garlic and onion adapted from Lanzotti et al. 2006 and Borlinghaus et al. 2019 (26, 47). LFS = Lachrymatory Factor Synthase.

Depletion of the glutathione pool through direct reaction between allicin and reduced glutathione is proposed as a major component of allicin’s antibacterial activity (9). We determined the percentage of total glutathione after 1 h of allicin treatment according to the procedure described by Müller *et al.* 2016 (9). PNA 97-1R WT maintained a higher percentage of glutathione compared to non-treated cells than a Δ*alt* strain indicating that the presence of the *alt* cluster counters allicin-mediated depletion of the glutathione pool (Fig. 3C).

We measured the total thiosulfinate content of full strength ROE to be 243 µM ±16 (n=8) using a 4-mercaptopyridine (4-MP) spectrophotometric assay (27). This is well above the allicin MIC values previously reported for *E. coli* (141.75 µM ± 10) and many other bacteria (7, 9). We also determined that allicin and onion thiosulfinates have similar capacity to delay the growth of PNA 97-1R and that the Δ*alt* mutant displayed consistently increased growth delay compared to WT PNA 97-1R across a range of allicin and onion thiosulfinate concentrations indicating similar antibacterial efficacy (Fig. 3D-E). We calculated the thiosulfinate MIC values of PNA 97-1R WT and Δ*alt* strains to be 125±5 µM and 80±5 µM respectively, based on unchanged OD_600_ at 24 hours in liquid LB incubated at 28°C.

We further monitored the contribution of *alt* to virulence based on systemic colonization of intact onion plants. This was conducted by tracking the capacity of Tn*7*Lux-labeled auto-bioluminescent PNA 97-1R derivative strains to systemically colonize onion bulbs 20 days after mechanical inoculation. The inoculations were conducted by piercing the onion neck just above the bulb shoulder. This form of inoculation is expected to mimic natural infection by simulating thrips-mediated feeding site preference and *Pantoea* transmission at the onion neck (16). The PNA 97-1R Δ*alt* strain showed a dramatic loss of auto-bioluminescence in the bulb at 20 DPI indicating reduced bacterial colonization (Fig. 4A). We observed a similar loss to bulb-associated autobioluminescence with a Δ*pepM* HiVir cluster mutant indicating that both loci are independently important for onion bulb colonization by PNA 97-1R (Fig. 4A). Similar patterns of auto-bioluminescence were seen during direct scale inoculation with Δ*alt* displaying the lowest bioluminescence. The Δ*pepM* strain, which critically does not induce necrosis associated with thiosulfinates release, showed higher scale colonization than the Δ*alt* strain, but limited to the zone around the inoculation puncture site. In contrast, the Δ*alt* Δ*pepM* strain showed less colonization than a Δ*pepM* strain presumably due to increased sensitivity to the thiosulfinate release associated with the inoculation puncture (Fig. 4B).

**Fig 4.**
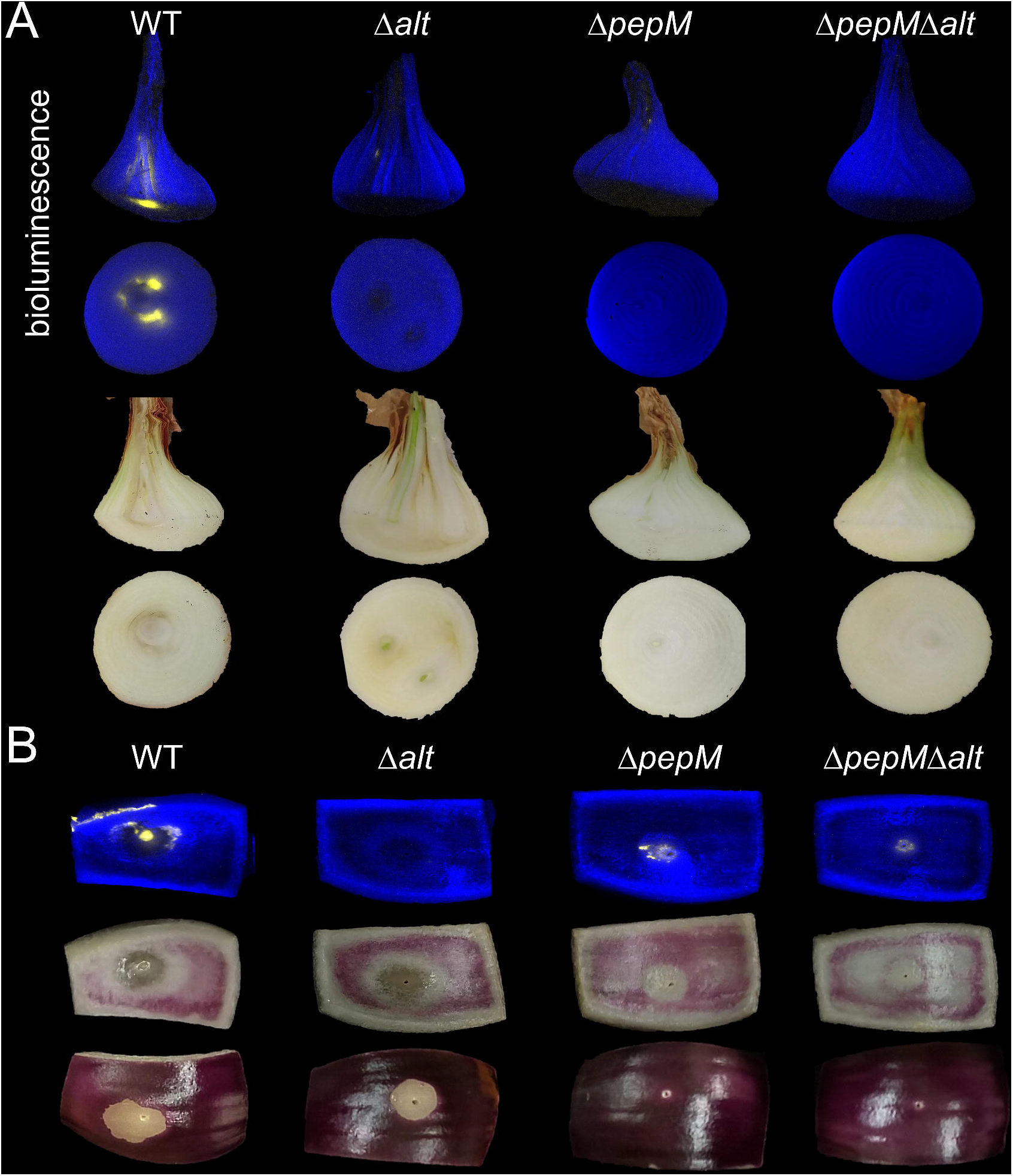
Onion bulb colonization of Tn*7*Lux labeled mutants 20 DPI. (*A*) Onions were inoculated at the neck with Tn*7*Lux labeled mutants 20 days prior to sampling. Onions were sliced longitudinal (top row) and transverse at bulb midline (bottom row). Bioluminescence was captured with 2 min exposure (yellow) and merged with the brightfield image (blue). Full color images were also taken of the same samples. Images are representative of three independent experiments and are not to scale. (*B*) Red onion scale colonization of Tn*7*Lux labeled mutants 3 DPI. Image signals are presented as described previously. Images are representative of three independent experiments and are not to scale.

### The *alt* cluster genes are sufficient for allicin tolerance and likely function as a cohort to tolerate thiol stress and promote onion scale colonization

We generated a series of nested complementation constructs carrying either the entire 11-gene *alt* cluster or sub-regions in the pBBR1-derivative plasmid pBS46 to determine their phenotypic contributions (Fig. 5A)(28). Complementation of PNA 97-1R Δ*alt* with the full *alt* cluster clone (*altB-J*) fully complemented onion scale colonization measured by both bacterial load and autobioluminescence, growth in ROE, and allicin tolerance based on both zone of inhibition and dilution plate assays (Fig. 5B-E, Fig. S5, S7-S10). Using the nested complementation constructs, the *alt* cluster sub-region *altB-G* displayed near wild type phenotypes in all assays. Sub-clusters *altB-A* and *altD-G* consistently showed partial complementation of *alt* phenotypes while *altH-altJ* showed minor independent phenotypic complementation in some assays. As these three *alt* sub-clusters share no genes in common, this supports a model of independent additive contributions of these genes to overall allicin tolerance and onion colonization phenotypes. The *altH-altI* genes, annotated as a carbon-sulfur lyase and cysteine-family-exporter genes, were not able to complement Δ*alt* phenotypes independently. The genes *altA* and *altR* have distant similarity to the *nemA* reductase and *nemR* TetR-family repressor genes of *E. coli* respectively that respond to bleach and N-ethylmaleimide and confer increased tolerance to these reactive compounds (29). A Δ*altR* mutant displayed reduced growth lag in ROE while *altR* overexpression complementation conversely displayed increased growth lag (Fig. S6). This is consistent with AltR serving to repress the ROE growth phenotype of the *alt* cluster.

**Fig 5.**
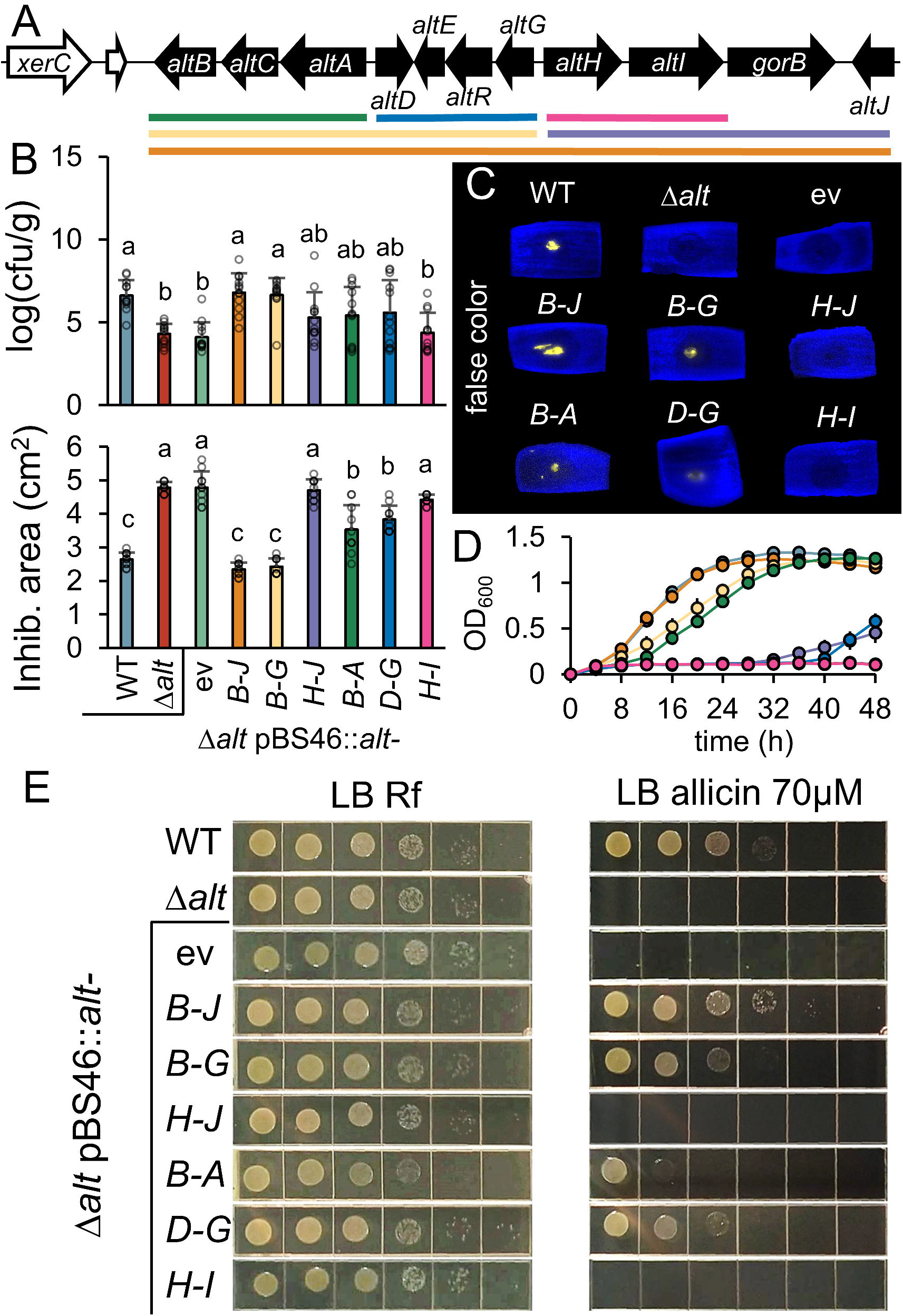
Nested complementation of the *alt* gene cluster. (*A*) Graphical representation of nested complementation clones spanning different regions of the *alt* cluster. Regions were cloned into the expression vector pBS46 and transformed into the PNA 97-1R ∆*alt* background. Colors indicate regions cloned and correspond to panels B and D. (*B*) Complementation clone bacterial load (top, *N=12*) and allicin tolerance (bottom, *N=9*) in the ∆*alt* background (ev = empty vector). The data from three independent replicates is presented. (one-way ANOVA followed by Tukey’s post-test, *p*<0.001, letters represent significant dif). Error bars represent ±SD. Log_10_ cfu/g, colony-forming units per gram of onion scale tissue. *(C*) Scale colonization of Tn*7*Lux labeled complementation clones 3 DPI. Bioluminescent strain signals were captured with 2 min exposure (yellow) and merged with the brightfield image (blue). Images are representative of three independent experiments. (*D*) Complementation constructs rescue growth in ROE (change in OD_600_). This experiment was repeated three times with similar results (*N=4*). Error bars represent ±SE. (*E*) Complementation clone ten-fold serial dilution of OD_600_ 0.3 on LB rifampicin and LB allicin amended plates Each row is a cropped from a larger photo representing one of three independent experimental replicates conducted in duplets. Original uncropped images (Fig S7-S10).

We expressed *altB-J* in the *P. ananatis* isolate PNA 02-18, which naturally lacks the OVR genes but possesses the HiVir gene cluster. The *altB-J* full cluster expression clone conferred multiple gain of function phenotypes on PNA 02-18 allowing the strain to colonize onion scales, grow well on ROE, and display increased tolerance to allicin (Fig. 6A-D, Fig. S5, S11). Transformation of *altB-J* into PNA 02-18 produced phenotypes consistent with complementation of the PNA 97-1R Δ*alt* strain. Similarly, heterologous expression of *altB-J* conferred increased allicin tolerance to *E. coli* DH5α indicating the *alt* genes are sufficient to confer allicin tolerance to bacteria outside of *Pantoea* (Fig. 6D-E, Fig. S12).

**Fig 6.**
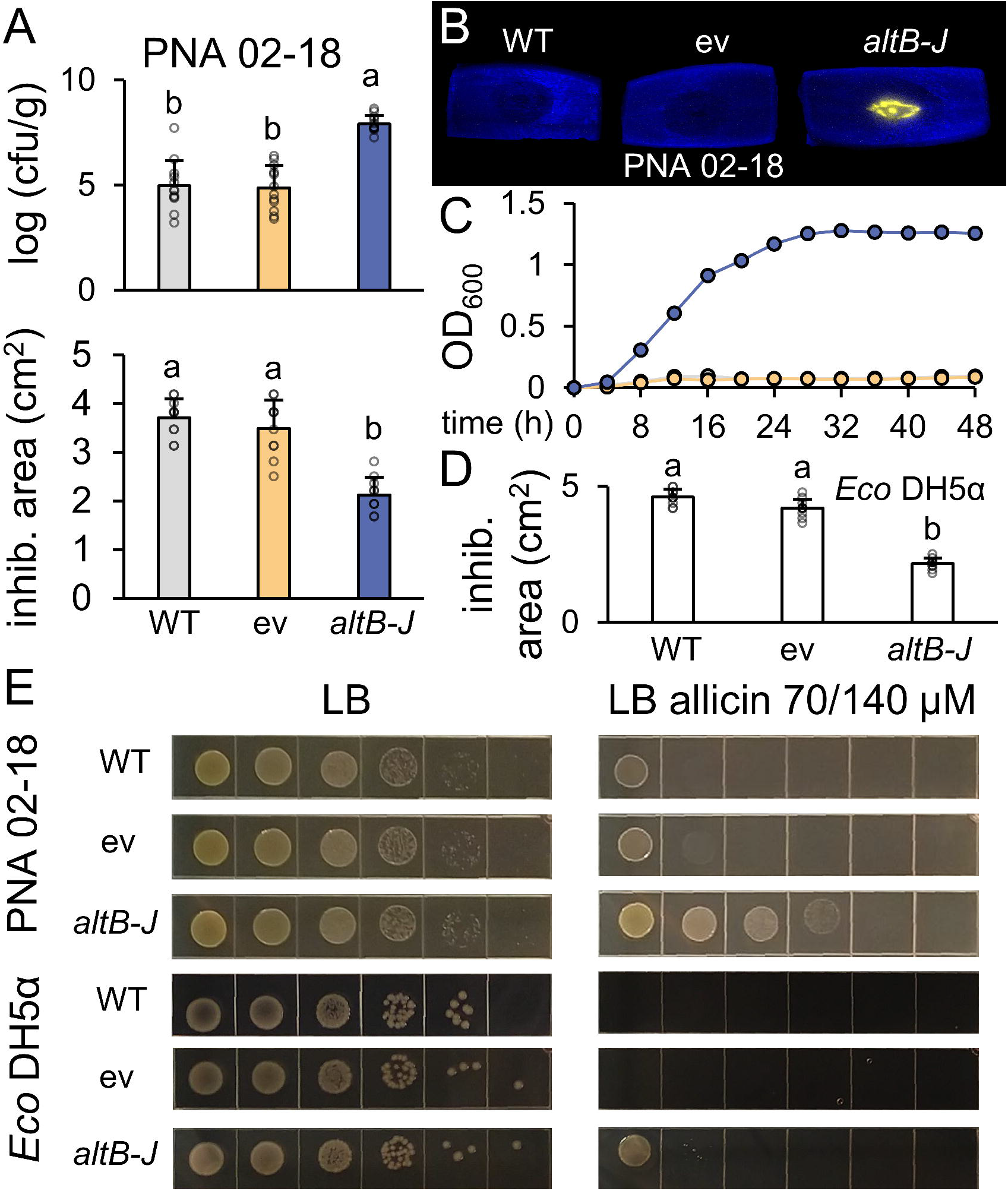
Complementation of *alt* associated phenotypes in heterologous backgrounds. (*A*) Bacterial load (top, *N=12*) and allicin tolerance (bottom, *N=9*) of heterologous expression clones in the PNA 02-18 naturally OVR-lacking background. The data from three independent replicates is presented. (one-way ANOVA followed by Tukey’s post-test, *p*<0.001, letters represent significant dif). Error bars represent ±SD. Log_10_ cfu/g, colony-forming units per gram of onion scale tissue. (*B*) Onion scale colonization of Tn*7*Lux labeled heterologous expression clones 3 DPI. Luminescent strain signals were captured with 2 min exposure (yellow) and merged with the brightfield image (blue). Images are representative of three independent experiments. (*C*) Bioscreen growth of PNA 02-18 *alt* expression clones in ROE over 48h. Error bars represent ±SE. This experiment was repeated three times with similar results (*N=4*). (*D*) Allicin tolerance of *E. coli* DH5α heterologous expression clones. The data from three independent replicates is presented (*N=9*, one-way ANOVA followed by Tukey’s post-test, *p*<0.001, letters represent significant dif). (*E*) Heterologous expression of *altB-J* in *P. ananatis* PNA 02-18 and *E. coli* DH5α in ten-fold serial dilution of OD_600_ 0.3 on LB and LB allicin amended plates. Each row is a cropped from a larger photo representing one of three independent experimental replicates conducted in duplets. Original uncropped images (Fig. S11, S12).

## Discussion

We observed a high degree of correlation between the genetic requirements for *P. ananatis* strains to colonize necrotized onion bulb tissue, grow in ROE, and their capacity for thiosulfinate tolerance. This suggests that tissue damage-induced endogenous production of thiosulfinates in onion exerts an antimicrobial effect on non-adapted bacterial strains and that thiosulfinates play an important role in biotic interactions between the onion host and bacterial pathogens.

The total thiosulfinate potential by *Alliums* may be an important factor both for disease outcomes and *Allium*-biotic interactions in general. Onion pathogenic *Burkholderia* have been shown to be sensitive to thiosulfinates *in vitro* as have many other plant pathogenic microbes (30, 31). In garlic, the level of host resistance to the chive gnat (*Bradysia odoriphaga*) was shown to correlate with cultivar-level variations in thiosulfinate production potential (32). *P. ananatis* routinely causes economically impactful outbreaks of center rot disease in sweet onions, which accumulate lower amounts of *S*-alk(en)yl-L-Cys sulfoxide precursor compounds (structures 1b, 1c) than other onions and thus, have comparatively less capacity for thiosulfinate production (5, 33). The presence of the *alt*-like cluster in onion-virulent *Enterobacter cloacea* EcWSU1 supports the idea that *alt* genes are adaptive for colonization of onion bulb tissue during disease. Interestingly, while there are many bacterial pathogens that cause disease in onion bulbs, there are comparatively few bacterial diseases of garlic bulbs (34). *Pseudomonas salomonii*, has been reported to cause “café au lait” disease on garlic leaves and carries gene clusters for allicin tolerance similar to those recently described in *P. fluorescens Pf*AR-1 (26, 35). Garlic leaves have less allicin production potnetial than garlic bulbs (36, 37). The garlic saprophytic *P. fluorescens* strain *Pf*AR-1 was found to carry three, nearly identical, chromosomal loci capable of conferring allicin tolerance with high specificity when heterologously expressed in *E. coli* or *Pseudomonas syringae* (26). However, while *Pf*AR-1 was isolated from garlic bulb, it is not a garlic pathogen and does not cause disease-associated necrosis on garlic. We speculate that three chromosomal allicin tolerance clusters of *Pf*AR-1 are potentially adaptive for stable saprophytic colonization of the garlic bulb niche. We presume that extreme thiosulfinate tolerance measures may be required to tolerate the potentially high thiosulfinate levels released via even minor coincidental wounding events over the life of the garlic bulb association.

The silencing of LFS has been used in laboratory settings to create “tearless” onions (5, 38). The lack of LFS activity in these lines drives increased flux of isoalliin into the dramatically increased production of 1-propenyl-based thiosulfinates after tissue damage (5, 38). Consistent with the capacity of garlic, which lacks an LFS-like enzyme, to produce nearly 100 fold more thiosulfinates than onion per g fresh weight, and the paucity of garlic bulb infecting bacterial pathogens in general, we hypothesize that these LFS-silenced onion lines should display increased resistance to *P. ananatis* infection by overwhelming the pathogen’s capacity for thiosulfinate tolerance. Conversely, onions with reduces alliinase activity rather than LFS activity would be expected to produce less thiosulfinates and have increased susceptibility to *P. ananatis* including naturally *alt*-lacking but HiVir containing isolates.

The ability of independent *alt* genes to confer partial phenotypic complementation and the functional predictions of the *alt* genes supports the model that the *alt* cluster encodes an additive cohort of proteins that collectively and cooperatively manage the impacts of cellular thiol stress as opposed to either direct inactivation or exclusion of thiosulfinates. Genetic dissection of the *Pf*AR-1 allicin tolerance clusters also demonstrated a cohort effect for allicin tolerance. The *Pf*AR-1 genes, *dsbA, trx*, and *aphD*, made the largest single gene contributions to allicin tolerance based on Tn insertional mutations and single gene overexpression tests (26). PNA 97-1R *alt* genes with similar annotations, *altC*, *altD*, and *altE* respectively, are all included on the *altB-G* sub-cluster clone, which conferred near wild type levels of complementation. The distant similarity of AltR to NemR could indicate a possible mode of action for AltR response to thiol stress. NemR carries a redox-sensitive cysteine residue critical for response to bleach and N-ethylmalimide and release of NemR from DNA to de-repress transcription of the NemA reductase (29). We hypothesize that thiosulfinate reaction with AltR cysteine residues may similarly allow AltR to respond to thiol stress, release from DNA and thereby de-repress expression of *alt* genes. Determining the roles played by specific Alt proteins in thiol stress perception and tolerance will be a fruitful area for future study.

In *P. ananatis*, the production of necrotic symptoms in onion, mediated by the HiVir chromosomal cluster, and colonization of necrotic onion tissue, mediated by the *alt* cluster, are genetically separable. The HiVir chromosomal cluster shows clear signs of horizontal gene transfer, while the plasmid-borne *alt* cluster is flanked by a recombinase-like gene (19, 20). Our disease model predicts that *P. ananatis* isolates lacking HiVir but possessing the *alt* cluster could benefit from co-associations with HiVir+, necrosis-inducing isolates as social cheaters. *P. ananatis* is a rare example of an aggressive gram negative plant pathogen that requires neither a T3SS to deliver virulence effectors to the plant cytosol nor a T2SS to deliver plant cell wall degrading enzymes to cause plant disease. This is unlike other well characterized examples of plant pathogenic *Pantoea*, *P. agglomerans* pvs. *betae* and *gypsophila* and *P. stewartii* subsp. *stewartii* that are dependent on a Hrp1-class T3SS for plant pathogenicity (39). Another characterized example of a Hrp-independent gram negative phytopathogen is *Xanthomonas albilineans* for which the phytotoxin albidicin, a DNA gyrase inhibitor synthesized by a hybrid PKS/NRPS pathway, is required to cause leaf scald disease of sugar cane (40, 41). Similarly, the ability of *P. ananatis* to cause necrosis on onion is dependent on the HiVir pathogenicity gene cluster, a non-PKS/NRPS gene cluster proposed to drive the synthesis of an, as of yet, undiscovered phosphonate secondary metabolite potentially functioning as a phytotoxin (20).

The pathogenic lifestyle of *P. ananatis* on onion more closely resembles that of some necrotrophic plant pathogenic fungi such as *Alternaria* or *Fusarium* than other described bacterial plant pathogens. These fungal pathogens produce phytotoxins as important virulence factors and proliferate on dead plant tissue (42, 43). In the case of *Fusarium oxysporum* f. sp. *lycopersici*, pathogen tomatinase enzyme-mediated resistance to antimicrobial tomatine saponins is critical for full virulence on tomato (44). Similarly, the *P. ananatis alt* genes confer tolerance to thiosulfinates and are required for colonization of necrotized onion tissue. We propose that a chemical arms race model provides a good framework for understanding *P. ananatis* disease on onion. *Pantoea* HiVir induces cell death in onion cells, potentially through synthesis of an undescribed phosphonate phytotoxin. Onion cell death coincides with post mortem generation of thiosulfinates which can restrict the growth of a non-adapted *P. ananatis*. The presence of the *alt* genes in *P. ananatis* confers tolerance to damage-induced onion chemical defenses and allows proliferation of adapted bacterial strains. This chemical arms race model for disease and defense in the *P. ananatis* onion pathosystem provides an interesting evolutionary contrast to the plant immune receptor/virulence protein effector arms race model that underlies many plant-pathogen interactions.

## Materials and Methods

### Bacterial strains and culture conditions

Overnight (O/N) cultures of *E. coli*, *Pantoea* spp., and *Enterobacter cloacae* were routinely cultured from single clones recovered on LB parent plates and were grown in 5 mL of LB media in 14 mL glass culture tubes at 28°C (*Pantoea*) or 37°C (*E. coli* and *E. cloacae*) with 200 rpm shaking (Table S6).

### Plasmid constructs and generation of mutants

Plasmids were typically constructed via Gateway cloning of PCR products, overlap-extension joined PCR fragments or synthesized DNA fragments. Deletion mutants were generated via allelic exchange using the pR6KT2G vector (*SI Appendix, Materials and Methods*, Table. S2-S5). Tn*7* transposants were generated essentially as described in Choi *et al.* 2008 (45). Expression plasmids were introduced via electrotransformation.

### Foliar necrosis assay

Onion seedlings (cv. ‘Century’, 12L:12D, six-weeks) were inoculated by depositing a 10 μl of bacterial suspension (OD_600_ 0.3 ≈ 1×10^8^ CFU/mL; dH_2_O) 1 cm from the central leaf apex on a wound created with sterile scissors. Leaves were evaluated 3 days post inoculation (DPI).

### Red onion scale necrosis assay

Consumer produce red onions were cut to approximately 3 cm wide scales, sterilized in a 3% household bleach solution for 1 m, promptly removed and rinsed in dH_2_O and dried. Scales were placed in a potting tray (27 × 52 cm) with pre-moistened paper towels (90 mL dH_2_0). Individual onion scales were wounded cleanly through the scale with a sterile 20 μl pipette tip and inoculated with 10 μl of bacterial O/N LB culture. Sterile deionized water was used as a negative control. The tray was covered with a plastic humidity dome and incubated at RT for 72 h. Following incubation, lesion sizes were measured by recording the diameter and small squares (0.06-0.08 g) of tissue were excised from a region 1 cm from the inoculation wound. Tissues were weighed and placed in plastic maceration tubes with beads, beat with a GenoGrinder SPEX SamplePrep 2010, ten-fold serially diluted with sterile dH_2_O and plated on rifampicin amended LB plates to determine the colony forming units per gram of onion tissue.

### Confocal imaging and staining

Scales were inoculated as described above. 100-125 mm^2^ sections were cut with a razor from the underside of onion scales and peeled at one corner with tweezers to minimize mechanical damage to other cells within the sample. Peeled samples were stained in fluorescein diacetate (FDA; 2 µg/mL) and propidium iodide (PI; 10µg/ml) at room temperature for 15 m in dark conditions as previously described (25). Stained samples were mounted on a slide in water under a coverslip for live-cell imaging. Confocal microscopy was performed with a Zeiss LSM 880 confocal microscope using the 10x objective.

Fluorescein was excited using 488 nm laser and emission collected between 508 and 535 nm. Z-stack imaging was used to image cells at multiple focal planes. PI was excited using a 543 nm laser and emission collected with 615-700 nm. Images were processed using the Zen software.

### Preparation of allicin and *alliaceous* extracts

Allicin was synthesized using the protocol of Albrect *et al.*, 2017 with some modifications (46). 15 µL of diallyl disulfide 96%, 25 µL of glacial acetic acid, and 15 µL of 30% H_2_O_2_ were added to a 200µL PCR tube and agitated for 4 h at 28°C. The reaction was quenched in 2mL of methanol. Fresh *alliaceous* extracts were prepared using the Breville Juice Fountain Elite or a kitchen blender. Solid debris were removed by straining subsequent macerates through cheese cloth and filter paper. Fine particulates were pelleted by centrifugation (10,000 g, 1.5 h, 4°C). Semi-clarified extracts were sterilized with a Nalgene 0.2 micron vacuum filter sterilization unit. All extracts were used within the week of preparation and stored at −20°C. Thiosulfinate concentrations were quantified using the 4-mercaptopyridine (4-MP) spectrophotometric assay (27).

### Liquid growth assays

Growth assays were conducted using 100-well honeycomb plates with the Bioscreen C system (Lab Systems Helsinki, Finland). The instrument was run for 48 h with low agitation at 28°C. Each well had 400 uL: 360 uL of the respective growth media and 40uL of an OD_600_ = 0.3 bacterial suspension in sterile dH_2_O, with a minimum of 3 well replicates. ROE was diluted with sterile water (≈ 100 µM thiosulfinate). ROE:LB media consisted of LB with an equal volume of ROE (≈ 80 µM thiosulfinate). Absorbance values were recorded every hour. Raw absorbance readings were normalized by subtracting the initial absorbance readings from subsequent hourly readings.

### Zone of inhibition assay

Styrene petri plates (100 × 15 mm) with 20 mL LB were spread with a bacterial suspension (OD_600_ 0.3 ≈ 1×10^8^ CFU/mL; dH_2_O) using a sterile cotton swab. 0.125 cm^2^ agar plugs were removed from the plates with a biopsy punch to create up to three wells per plate. 50 µL of either garlic extract or allicin stock solution were added to the wells. Plates were incubated for 24 h at RT and evaluated for a zone of inhibition (cm^2^) by measuring the radius, calculating the inhibition, area and subtracting the well area.

### Serial dilution allicin senistivity plates

The allicin stock solution was added to molten LB cooled to 55°C in sterile conical vial tubes and poured into square plates to set, achieving relative allicin concentrations of 70 µM for *Pantoea* and 140 µM for *E. coli*. The O/N cultures of strains to be tested underwent ten-fold series dilutions and 10 uL volumes of each dilution were plated on the LB control and allicin-amended LB plates. The plates were incubated overnight (28°C *Pantoea*, 37°C *E. coli*) and imaged the following morning.

### Sweet onion systemic infection

Sterile wooden toothpicks were soaked in a bacterial suspension (OD_600_ 0.3 ≈ 1×10^8^ CFU/mL; dH_2_O) and were inserted horizontally through the entire neck of five-month-old the onion plants (cv. Sapelo Sweet) just below the leaf fan and above the shoulder. Toothpicks were left in the plants. At 20 d the onions were harvested for imaging. Onions were removed from soil (which was sterilized following the experiment and discarded), rinsed with water, and cut twice horizontally at the center of the bulb to produce a 1.5 cm section of the center of the onion. The remaining portion of the top of the bulb had foliage removed and was cut vertically. Sliced onions were imaged with a color camera followed by long exposure imaging with the analyticJena UVP Chemstudio.

### Quantification of glutathione

Glutathione quantification was conducted essentially as described by Müller *et al.* 2016 using a Glutathione (GSH) Colorimetric Detection Kit (Arbor Assays) (9). Optical readings were conducted with Tecan Spectra Rainbow spectrophotometer in 96 well styrene plates and the glutathione concentration of the samples were determined according to the manufacturers recommendations.

### False color luminescence images

Luminescence false color images were merged in ImageJ to create compound images of luminescence and brightfield captures. The TIFF file of the long exposure capture (luminescence) and brightfield capture were opened with ImageJ Fiji release (https://imagej.net/Citing). The images were merged (image→ mergechannels→ [select blue for brightfield and yellow for luminescence]). The output file was saved in PNG format for publication images.

## Acknowledgements

We would like to acknowledge Nathan Weinmeister for assistance with allelic exchange to create the pepM/*alt* double mutant, Brenda Schroeder and Cheryl Patten for providing *Enterobacter* strains, Ron Walcott, Marin Brewer, and Li Yang for use of equipment, Paul Severns for statistical consulting, and members of the Yang and Kvitko Lab for helpful discussions regarding the preparation of the manuscript. We acknowledge the assistance of the Biomedical Microscopy Core at the University of Georgia with imaging using a Zeiss LSM 880 confocal microscope. This work was supported with funding from the Vidalia Onion Committee, USDA SCBGP Project AM180100XXXXG014, and USDA NIFA SCRI Project 2019-51181-30013.

---

Fig S1. Growth of natural variant *Pantoea* isolates in aqueous red onion extract (ROE) and ROE amended with LB. (*A*) Growth of isolates (change in OD_600_) in ROE. (*B*) Growth of isolates in ROE:LB. This experiment was repeated three times with similar results (*N=6*). Error bars represent ±SE.

Fig S2. Growth of PNA 97-1R ∆*alt* in LB broth amended with allicin. Bioscreen growth of strains (change in OD_600_), PNA 97-1R WT and PNA 97-1R ∆alt. (*A*) synthesized allicin amended LB. (*B*) garlic extract amended LB. Allicin concentration determined with 4-MP assay. This experiment was repeated three times with similar results (N=4). Error bars represent ±SE.

Fig S3. PNA 97-1R *alt* cluster shares homology with a plasmid-borne gene cluster of onion rot pathogen *Enterobacter cloacae* EcWSU1. Percent amino acid identity is depicted above homologous EcWSU1 genes. NCBI locus numbers are included for EcWSU1 genes.

Fig S4. Natural variants of *P. ananatis* and *Enterobacter cloacae* with *alt*-like genes have increased tolerance to allicin and garlic extracts. Area of inhibition for garlic extract (top) and allicin (bottom) is represented. Representative zones of inhibition for allicin above corresponding bars. The data from one of three independent replicates is presented (*N=6*, one-way ANOVA followed by Tukey’s post-test) (*N=6*, t-test, ***p<0.0001). Error bars represent ±SD.

Fig S5. Calculated total lesion luminescence (CTLL) of complementation constructs in WT and heterologous *Pantoea* background. Values were quantified by defining the lesion area in stacked images and quantifying relative pixel intensity in imageJ. CTLL= integrated density – (area of lesion X mean luminescence of background readings, ctrl). (*A*) CTLL of PNA 97-1R nested complementation constructs. The data from six independent replicates is presented (*N=18*, one-way ANOVA followed by Tukey’s post-test). (*B*) CTLL of PNA 02-18 expressing *altB-J.* The data from three independent replicates is presented (*N=12*, one-way ANOVA followed by Tukey’s post-test).

Fig S6. Growth curve of *altR* mutants in LB:ROE. Bioscreen growth of isolates (change in OD_600_), ev = empty vector. This experiment was repeated three times with similar results (*N=6*). Error bars represent ±SE.

Fig. S7. WT, ∆*alt*, *altH-J* ten-fold serial dilution on LB rifampicin and LB allicin amended plates. Fig. 5 cropped panels highlighted in red.

Fig S8. WT, ∆*alt*, *altH-J* ten-fold serial dilution on LB rifampicin and LB allicin amended plates. Fig. 5 cropped panels highlighted in red.

Fig S9. WT, ∆*alt*, *altH-J* ten-fold serial dilution on LB rifampicin and LB allicin amended plates. Fig. 5 cropped panels highlighted in red.

Fig S10. WT, ∆*alt*, *altH-J* ten-fold serial dilution on LB rifampicin and LB allicin amended plates. Fig. 5 cropped panels highlighted in red.

Fig S11. *P. ananatis* PNA 02-18 WT, ev, and *altB-J* ten-fold serial dilution on LB and LB allicin amended plates. Fig. 6 cropped panels highlighted in red.

Fig S12. *E.coli* DH5α WT, ev, and *altB-J* ten-fold serial dilution on LB and LB allicin amended plates. Fig. 6 cropped panels highlighted in red.

