## Supplemental data and methods for "*Pantoea ananatis* defeats *Allium* chemical defenses with a plasmid-borne virulence gene cluster"

**Supporting Information Appendix**

### Materials and Methods

#### Bacterial strains and culture conditions.

Overnight (O/N) cultures of *E. coli, Pantoea,* and *Enterobacter cloacae* were routinely cultured from single clones recovered on LB parent plates and were grown in 5 mL of LB media in 14 mL glass culture tubes at 28°C (*Pantoea*) or 37°C (*E. coli* and *E. cloacae*) with 200 rpm shaking. When required antibiotics and media additives were used at these concentrations in liquid and solid culture: 50 µg/mL spectinomycin (Sp), 10 µg/mL gentamicin (Gm), 25 µg/mL chloramphenicol (Cam), 50 µg/mL kanamycin (Km), 60 µg/mL rifampicin (Rf), 60 µg/mL 5-bromo-4-chloro-3-indolyl-beta-D-glucuronic acid (X-Gluc), 200-400 µg/mL diaminopimelic acid (DAP).

#### General.

*PCR.* Primers were ordered from Eurofins Genomics USA (Table. S2, Table. S3). Genomic DNA was routinely extracted using the Gentra Puregene Yeast/Bact. Kit (Qiagen). DNA was quantified with the Eppendorf Biospectrometer basic. PCR reactions used the analyticjena FlexCycler2 with the following protocol for 50 μl reactions: 20 μl 5X HiFi buffer, 5 μl forward primer, 5 μl reverse primer, 2 μl 10 mM dNTPs, 1 μl genomic DNA, 1 μl Phusion High Fidelity Polymerase (New England Biolabs), 66 μl PCR grade dH_2_O under the following conditions 2 m @ 98°C; repeat 30 cycles, 10 s @ 98°C, 30 s @ 50°C, 30 s/kb @ 72°C, 10 m @ 72°C, Hold @ 4°C. Nucleic acids were routinely purified in 1.5% TBE gels or directly from reactions using Monarch PCR & DNA Cleanup Kit (NEB) or Monarch DNA Gel Extraction Kit (NEB).

*Restriction cloning.* Restriction enzymes and T4 DNA ligase were ordered from either NEB or ThermoScientific and reactions were completed by following manufacturer protocols or the NEBcloner tool (v1.3.14). Plasmids were extracted with the GeneJet Plasmid MinPrep Kit (ThermoScientific). Restrictions were simulated using NEBcutter V2.0. Vector constructs were sequenced using Eurofins Genomics with either M13 standard primers or self-designed sequencing primers and confirmed by mapping to simulated vectors created in the Geneious application (Geneious v2019.1.3) (Table. S2, Table. S3)

*Gateway cloning*. Gateway cloning included BP clonase reactions into the pDONR221 cloning vector and pR6KT2G suicide vector and subsequent LR clonase reactions into the pBS46 expression vector. Reactions followed manufactures protocols (Thermoscientific).

*Computational programs*. Calculations, graphs and figures were made using the Microsoft Office 365 programs Excel, Power Point, and Word. Basic genomic sequence analysis including alignments, primer design, organizing sequencing data, vector construction, and reaction simulations were conducted using Geneious Prime v2019.1.3. Statistical analyses including Tukey-HSD ANOVA and student t-test were completed in R-Studio v1.2.1335 (package agricolae). Image analysis and manipulation was conducted using ImageJ (Fiji release).

#### Maintenance of onions.

*Foliar onion seedling assay*. Onion seedlings (*Allium cepa* L. cv. ‘century’; six weeks) were established in 10 cm × 8 cm (diameter × height) plastic pots containing commercial potting mix. The seedlings were kept in a greenhouse and maintained at 25-28°C, 80-90% relative humidity, and 12L:12D photoperiod.

*Onion greenhouse experiment*. Onion sets (cv. Sapelo Sweet) were obtained from the Vidalia Onion and Vegetable experiment station (Lyons, GA) and were potted in 16 cm × 15 cm (diameter × height) plastic pots with commercial potting mix on Jan 3^rd^ and 4^th^ 2019. The onions were grown until May 8^th^, 2019 when they were inoculated.

*Red onion scale assays.* Large red onions were purchased from commercial grocery stores with typical weight ranging from 400-600g.

#### Construction of pR6KT2G suicide vector.

pR6KT2G was derived from pR6KT2 (Table. S2) (1). We sought create a Gateway compatible version of the pR6KT2 suicide vector to increase construction efficiency. Primers with NotI digest overhangs and TAAT extensions were designed to amplify the Gateway cassette of pDONR221 (Table. S3). The predicted product size was 2.5 kb and carried the cat promoter, Cm resistance cassette and bacterial gyrase toxin CcdB. Primers amplified the cassette and the insert was purified and eluted to 10 μl, digested with NotI, purified again, and ligated into the NotI digested pR6KT2 suicide vector. Drop dialysis of ligation reaction was conducted by placing the reaction volume on a VMWP membrane (Millipore) suspended on sterile dH_2_O for 30 m. The resulting reaction volume was electrotransformed into *Eco* DB3.1 *pir^+^* strains and recovered on Gm-Cam LB plates. Recovered clones were patched onto Km plates to ensure pDONR221 was not carried over. Putative clones were plasmid prepped, screened via BsrGI digest, and sequence confirmed.

#### Construction of the pTn7PA143LuxFK vector.

The synthetic bacterial promoter PA1/04/03 was synthesized as a dsDNA gblock by IDT (sequence below) and cloned into the DraIII digested backbone of pTn5/7LuxK6 (2) via Gibson assembly. The insertion of the PA1/04/03 promoter was confirmed by sequencing. TGGAACTAGATTTCACTTATCTGGTTGGCCTGCAAGGCCTCACtatGTGAaagagtgttgacttgtgagcggataacaatgatacttagattcaatCACgcgGTGacaagtttgtacaaaaaagcaggctgcgGTGCATTAAATGGATGGCAAATATGA

#### Closure of PNA 97-1R genome.

PNA 97-1R Genomic DNA was sequenced previously via Illumina sequencing by MicrobesNG (3). This workflow generated a total of 354,376 trimmed paired reads and 144,592,000bp of sequence (∼29× coverage).

A separate isolation of genomic DNA was extracted using a Nanobind kit (Circulomics, Baltimore MD) and was sequenced by the Baltrus lab via an Oxford Nanopore MinION device. 500ng of DNA barcoded using the prepared using barcode 1 of the native barcode expansion kit followed by preparation of fragments for sequencing using the LSK-109 kit without shearing. Reads were demultiplexed with qcat version 1.0.7. Reads were called during sequencing using Guppy version 2.0.10, and only the reads which passed the initial quality check (QC) during the run were used. Sequencing on the MinION device generated 160,932 reads, with a read distribution N50 of 10,336bp, for a total of 271,539,000bp of sequence (∼54× coverage).

Hybrid assembly of all read types was performed using Unicycler version 0.4.8 (4) and resulted in a single chromosome of 4,558,720bp of sequence with a 53.6% GC content and two complete plasmids (273,809bp and 161,611bp). Unicycler also specified that this chromosomal assembly was circular. This chromosomal sequence was annotated by the NCBI Prokaryotic Genome Annotation Pipeline (PGAP) version 4.8 (5) and it is predicted to contain 4,731 genes representing 4,616 predicted protein coding sequences, 7 complete sets of rRNAs (5S, 16S, and 23S rRNAs), 78 tRNAs, and 15 noncoding RNAs (ncRNAs). Default parameters were used for all software.

#### Creation of mutant strains by allelic exchange.

*Unmarked deletion. Pantoea ananatis* knockout clones were generated in the PNA 97-1R background using the pR6KT2G suicide vector which allows for SacB mediated sucrose counter-selection. Primers were designed to amplify flanks preceding (flank A ≈ 700bp) and following (flank B ≈ 800bp) the deletion target (Table. S3). Flank A-forward and flank B-reverse primers had attb1&2 5’ extension sites appended. Flank A-reverse and flank B-forward primers had overlap extension 5’ tails appended (Table. S3). Flanks A and B were amplified separately, purified, and joined via overlap-extension PCR in a 50 μl reaction under the following protocol: flanks were diluted ten-fold and master mix A was prepared: 5 μl HF 5X Buffer, 2 μl dNTPs, 0.25 μl Phusion polymerase 12.75 μl dH20 PCR water, 2.5 μl ten-fold dilution of flank A, 2.5 μl ten-fold dilution of flank B. Master mix B was prepared: 5 μl HiFi buffer, 2 μl 2.5mM dNTPs, 0.25 μl Phusion polymerase, 5 μl Flank A forward primer, 5 μl Flank B reverse primer, 7.75 μl of sterile miliQ water. Master mix A was added to a 200 μl PCR reaction tube under the following conditions: 5 m @ 94°C; repeat 10 cycles, 30 s @ 94ºC, 1.5 m @ 60°C, 2.5 m @ 72°C, 10 m @ 10°C (add Mix B), repeat cycle 35 times, 30 s @ 94°C, 30 s @ 58°C, 2 m @72°C, 10 m @72°C, hold 4°C. The product was run on a 1% agarose gel stained with ethidium bromide to visualize the fusion product for gel extraction (A+B ≈ 1,500bp). The fusion band was excised, and gateway cloned into pR6KT2G [if over-lap extension PCR was unsuccessful deletion double stranded DNA gblocks were synthesized by Integrated DNA Technologies]. Drop dialysis of the BP clonase reaction was conducted as previously reported. The reaction mixture was electrotransformed into *Eco* MaH1 *pir^+^* strains and recovered on Gm plates. Six clones recovered from the transformation were grown O/N, plasmid prepped, digest confirmed with XhoI, and sequence confirmed. Confirmed deletion constructs were then transformed into E-comp *Eco* RHO5 *pir*^+^ mating strain. The wild-type (PNA 97-1R) and donor strain (RHO5 [pR6KT2G_XX]) were mated on DAP amended LB plates O/N. The following day merodiploid were recovered on Gm plates. Two merodiploid colonies were selected to inoculate a liquid culture comprised of 1 mL sterile LB and 3 mL sterile 1M sucrose. The counter selection culture was incubated at 37°C for 24 h. Following counter selection 200 μl of a 1×10^6^ dilution of the counter selection culture was plated on X-gluc amended LB plates. Colonies that were yellow had evicted the plasmid construct. Putative mutants were genome prepped and screened using “out” primers (Table. S3). PCR positive clones were then sequenced to confirm the deletion of the gene or gene cluster in question.

*Marked deletion.* In the case of generating marked deletions (see altR mutant) the Sp resistance cassette from pCPP5242 was cloned between the two deletion flanks (Table. S2, S3). The confirmed suicide vector was digested with AvrII and the Sp resistance insert was digested with NheI. The products were ligated, transformed, plasmid prepped, digest confirmed, sequence confirmed, transformed into *Eco* RHO5 *pir*^+^, mated with target strain, and two merodiploid clones recovered on Gm plates. The same sucrose counter selection was employed as described above but the resulting culture was diluted and plated on LB Sp plates instead of x-gluc amended LB. Putative mutants were genome prepped and screened with “out” primers (Table. S3).

#### Mini-Tn*7*Lux labeling of strains.

*Eco* RHO3 pTNS3, *Eco* RHO5 pTn*7*PA143LuxFK, and the target *Pantoea* strain were combined in a tri-parental mating (Table. S2, S6) (6). 5 mL LB cultures of each strain were grown O/N. The following day 1 mL of each culture was pelleted and resuspended in 100 µL of fresh LB. 20 µL of each concentrated culture was added to a clean tube. DAP amended LB plates were prepared and sterile nitrocellulose membranes were placed on the plates. 20-30µL droplets of the mixed cultures, and independent strain controls were placed on the nitrocellulose membranes to dry. Following O/N incubation, the mixture was removed from the nitrocellulose membrane using a sterile loop and resuspended in 1 mL LB. 100 µL of the incubated mixed culture suspension was plated on Km LB selection plates. The remaining volume was pelleted, resuspended, and plated. The following day Km resistant colonies were selected confirmed for luminescence with 2 m exposure settings on the lab imager (analyticJena UVChemStudio).

#### Foliar necrosis assay.

Seedlings were inoculated by cutting the central leaf 1 cm from the apex with a sterile pair of scissors. Using a micropipette a 10 μl drop of a bacterial suspension in sterile dH_2_O adjusted to OD_600_ 0.3 ≈ 1×10^8^ CFU/mL was placed at the cut end of the seedlings. Seedlings inoculated with dH_2_O were used as negative controls.

#### Red onion scale necrosis assay.

*Necrosis assay.* Consumer produce red onions (*Allium cepa*. L., red onion) were purchased, cut to approximately 3 cm wide scales, sterilized in a 3% household bleach solution for 1 m, promptly removed and rinsed in dH_2_O. Scales with a healthy unmarred appearance were used. Scales were placed in a potting tray (27 × 52 cm) containing two layers of paper towels pre-moistened with 90mL of distilled water. The plastic removable portion of 20 μl pipette trays were positioned on top to prevent direct contact between the paper towels and the scales. Individual onion scales were wounded cleanly through the scale with a sterile 20 μl pipette tip and inoculated with a 10 μl drop of bacterial O/N LB culture. Sterile deionized water was used as a negative control. The tray was covered with a plastic humidity dome and incubated at RT for 72 h. Measurements of the necrotic tissue area (radius1 × radius2) were recorded and the necrotic lesion area was calculated (r1×r2 × π).

*Quantification bacteria (CFU/g) within onion scales.* Following the 72 h incubation of inoculated scales, small squares (0.06-0.08 g) of tissue were excised from a region 1 cm above the inoculation wound. The excised tissues were weighed and placed in plastic maceration tubes with 100 μl of sterile dH_2_O and three polyethylene beads. The tubes beat at 1750 (strokes / m) for 1 m (GenoGrinder SPEX SamplePrep 2010). The macerate underwent a ten-fold series dilution with sterile dH_2_O in 96-well styrene plates (20 μl, 180 μl). Following the dilution 10 μl of the diluent was plated on Rf LB square plates and incubated for 24 h 28°C. Visible colonies were counted and the CFU concentration per gram was back calculated for each individual scale tested.

*Imaging of Tn7Lux labeled strain scale colonization.* Once inoculated with a Tn*7*Lux labeled strain and 72 h incubation, scales were removed and imaged with the analyticJena UVP Chemstudio. Within the visionworks software manual long-exposure imaging was selected with the following settings: capture time 2 m, 70% focus, and 100% brightness(aperture), stack image and saved to TIFF format. Brightfield images were captured with the following settings: capture time 20 mS 70% focus, 70% brightness(aperture), and saved to TIFF format.

#### Confocal imaging and staining.

*Onion Scale Peels. Pantoea ananatis* was used to infect onion scales as described above. Onion scales were subsequently incubated at room temperature in darkness for five to seven days. 100-125 mm^2^ sections were cut with a razor from the underside of onion scales and peeled at one corner with tweezers to minimize mechanical damage to other cells within the sample.

*Cell Staining.* The peeled samples were stained in fluorescein diacetate (FDA; 2 µg/mL) and propidium iodide (PI; 10µg/ml) at room temperature for 15 minutes in dark conditions as previously described (Jones et al., 2016). Stained samples were mounted on a slide in water under a coverslip for live-cell imaging.

*Microscopy.* Confocal microscopy was performed with a Zeiss LSM 880 confocal microscope using the 10x objective. Fluorescein was excited using 488 nm laser and emission collected between 508 and 535 nm. PI was excited using a 543 nm laser and emission collected with 615-700 nm. Images were processed using the Zen software (Black edition, version 2.3, Zeiss). Z-stack imaging was used to image cells at multiple focal planes.

*Image Analysis.* Images were taken near the center of onion peels to reduce imaging cells near mechanical damage. Live and dead cells were manually counted by observing z-stack images from uninfected and infected regions of stained onion peels. Live cells include those with typical FDA (green) staining of the cytoplasm and nucleus or without PI (red) stained nuclei. Dead cells include those with PI (red) stained nuclei, as previously described (Jones et al., 2016). Five separate images collected from different uninfected and infected regions of onion scales were used to quantify live and dead cell numbers, respectively.

#### Preparation of onion and garlic extracts.

*Blender method – aqueous extract.* Garlic (four bulbs) or onion (two large bulbs) were selected from local grocery establishments. The onions or garlic were sliced and blended with 300 mL of dH_2_O in a kitchen blender. The resulting macerate was filtered through wire mesh to remove debris. The filtrate was centrifuged in a 50 mL plastic conical vial tube (3900 g, 20 m, 4°C). The supernatant was carefully removed and filtered through cheese cloth and several filter papers (Whatman 1001-185). Finally, the aqueous extract was processed through a Nalgene disposable vacuum filter sterilization unit. The aqueous extract was either used that week or frozen in a -20C freezer for use in the future. Each time the mixture was used the allicin content was quantified using the 4-MP assay.

*Juicer method- crude extract.* One garlic bulb (60-80 g) or red onion bulb (400-600 g) was run through an industrial strength juicer (Breville Juice Fountain Elite) yielding 20-30 mL of crude garlic extract or 300-400 mL of crude red onion extract (ROE). The crude extract was then spun in a 250 mL plastic centrifuged (10,000 g, 1.5 h, 4°C). Following centrifugation, the upper liquid phase was removed carefully with a pipette and processed through a Nalgene disposable vacuum filter sterilization unit

#### Preparation of allicin stock solution.

Allicin is a relatively unstable molecule, thus fresh stock solutions must be prepared for experiments utilizing this chemical. Allicin was synthesized using a modified protocol from Albrecth et al. 2017 (7) 5 µL of diallyl disulfide 96% (Carbosynth), 25 µL of glacial acetic acid (Sigma aldrich), and 15µL of 30% H_2_O_2_ were added to a 200µL PCR tube. The PCR tube was taped to a 1.5 mL Eppendorf tube willed with water. The taped tubes were suspended together in a 500 mL glass beaker, and placed in a 28°C shaker for 4 hrs. The suspended contraption allowed for constant high-speed agitation of the small tube, facilitating the reaction. Following agitation, the reaction was quenched in 2mL of methanol. This methanolic allicin mixture was used as the stock synthesized allicin preparation and was used in extract inhibition assays. A negative control mock preparation lacking diallyl disulfide was included in inhibition assays for comparison.

#### Quantification of thiosulfinates (allicin).

Allicin and onion thiosulfinate concentrations were quantified using a spectrophotometric assay described by Miron et al. 2002 (8). The relative concentration of thiosulfinates in a sample is determined based on a negative optical absorbance shift that occurs when 4-mercaptopyridine (4-MP, λ_max_= 324 nm) forms a disulfide bond with the active thiosulfinate, producing 4-allylmercaptothiopyridine, which has no absorbance at 324 nm (8). The thiosulfinate activity is can be quenched by chemicals present in complex media. To account for this the thiosulfinate concentration of LB:(ROE; garlic; allicin) was measured 15 m after mixing the solutions. A dilution series of the extracts in LB was made and 50 µL of each dilution with a negative LB control was added to individual 950 µL aliquots of (1x10^-4^ M) 4-mercaptopyridine 50mM Na-phosphate, 2mM EDTA, pH 7.2. The reactions were gently agitated, incubated at RT 30 m, and ∆*A*_324_ was recorded_­_ in 2 mL styrene cuvettes. Absorbance values (∆*A*_324_) between 0.08-0.3 gave the most consistent result using the Eppendorf Biospectrometer basic. Values within this range were used to back calculate stock concentrations from series dilutions. The following equation was used to convert the absorbance values into the molar concentration of thiosulfinates: [allicin] = ∆*A_324_* × 50[dilution])/39,600 M (8). The average stock concentration of freshly prepared ROE, garlic, and synthesized allicin were ≈ 250.37±26.7 µM, 7621.6±527 µM, and 9242±200 µM respectively.

#### Strain growth in Bioscreen C.

The absorbance value OD_600_ was used extensively to represent bacterial growth in media amended with various extracts. The growth assays were conducted using honeycomb plates in the Bioscreen C (Oy Growth Curves Ab Ltd). The instrument was run for 48 h with low agitation in 400 uL wells with 360 uL of the respective growth media and 40uL of an OD_600_ = 0.3 bacterial suspension in sterile dH_2_O. The absorbance value was recorded every hour over this time period. The raw bioscreen absorbance readings were normalized by subtracting the initial absorbance readings from subsequent hourly readings. Each assay was typically conducted with at least four technical replicates and was repeated in its entirety three times.

#### Zone of inhibition assay.

Styrene plastic petri plates (100 × 15 mm) were filled with 20 mL of melted LB. A sterile cotton swab was wetted with an OD_600_ 0.3 ≈ 1×10^8^ CFU/mL suspension of the bacteria in sterile dH_2_O. The back end of a 200 µL pipette tip was used to create a 0.125 cm^2^ well in the petri dish, with up to three wells per plate. 50 µL of either garlic extract or allicin stock solution was added to the well. After incubation for 24 h the plate was evaluated for a zone of inhibition (cm^2^) by measuring the radius, calculating the inhibition, area and subtracting the well area.

#### In-vivo quantification of *Pantoea* glutathione.

*Pantoea ananatis* PNA 97-1 WT and PNA 97-1 ∆*alt* were incubated in 5 mL of LB O/N. 2.5 mL of this culture was used the following day to seed a 50 mL LB culture that was incubated O/N in a sealed plastic conical vial tube. The following morning 1 mL of stock allicin solution was added to the tubes for a molar concentration of 230 µM, an untreated sample was used as a control. The tubes were incubated for 30 m and then spun at 4,000 g for 10 min. The supernatant was poured off and pelleted bacteria were resuspended in 1 mL of KPE buffer. The cells were washed twice with fresh KPE and resuspended in a 5% (w/v) sulfosalicylic acid solution for deproteination. The cells were lysed with freeze thaw cycling (freeze -80 10 m, thaw 40° C bath 1 m 3X). Lysed cells were centrifuged at 10,000 g for 5 min. 50 uL of deproteinated samples were used in accordance with the Glutathione (GSH) Colorimetric Detection Kit (Arbor Assays). Optical readings were conducted with Tecan Spectra Rainbow spectrophotometer in 96 well styrene plates and the Glutathione concentration of the samples was determined using the manufacturers myassays online tool. The assay was used to determine total glutathione in bacteria after allicin treatment and total glutathione in the untreated control samples. The percent change from the untreated control was calculated by dividing the allicin treated glutathione concentration by the paired control glutathione concentration and multiplying by 100.

#### Sweet onion neck stab assay and imaging.

*Inoculum preparation.* Cultures were grown O/N in LB with selective antibiotics. The inoculum concentration was adjusted to OD_600_ 0.3 ≈ 1×10^8^ CFU/mL in sterile dH_2_O with final a final volume of 25mL. Sterile toothpicks were soaked in the inoculum suspension for 20 m.

*Experimental design.* Colored tags were used to mark treatments. Each treatment was presented once in a block. Blocks were randomized and four blocks were included at each time point (20 d, 30 d, 40d).

*Inoculation*. To inoculate plants sterile toothpicks were removed from the bacterial suspension and stabbed horizontally through the entire neck of the onion plant just below the leaf fan. Toothpicks were left in the plants. At 20 d May 28^th^ , 30 d June 6^th^ , 40 d June 17^th^ the onions were harvested for imaging. The onions were removed from soil (which was sterilized following the experiment and discarded), rinsed with water, and cut twice transversely at the center of the bulb to produce a 1.5 cm section of the center of the onion. The remaining portion of the top of the bulb had foliage removed and was cut longitudinally.

*Imaging.* Sliced onions were imaged with a color camera followed by long exposure imaging with the analyticJena UVP Chemstudio. Within the visionworks software manual long-exposure imaging was selected with the following settings: capture time 2 m, 70% focus, and 100% brightness(aperture), stack image and saved to TIFF format. Brightfield images were captured with the following settings: capture time 20 mS 70% focus, 70% brightness(aperture), and saved to TIFF format.

#### Serial dilution allicin plate assay.

The allicin stock solution was added to melted LB in conical vial tubes and poured into square plates to set achieving relative allicin concentrations of 70 µM for *Pantoea* and 140 µM for *E. coli*. O/N cultures of strains to be tested underwent ten-fold series dilutions and 10 uL droplets of each dilution were plated on the LB control and LB allicin plates. The plates were incubated overnight (28°C *Pantoea* 37°C *Eco*) and imaged the following morning.

#### Quantification of lesion luminescence.

Calculated lesion luminesce was conducted using the imageJ Figi release. The brightfield and luminesce TIFF grayscale images were opened in the program stacked (image→ stacks → images to stack). The circle selector was used to create a selection highlighting the onion lesion. The scroll was used to move from the brightfield layer to luminesce layer while maintaining the lesion area selection. The integrated intensity and mean grey values were quantified for the selected area (CTRL+M). Three negative regions were highlighted and measured for background control readings. The corrected total lesion luminescence was calculated with the following equation CTLL = integrated density – (area of selected lesion × mean grey value of background readings).

#### *Pantoea* - *Enterobacter* sequence homology analysis.

Geneious Prime V 2019.1.3 was used to convert nucleotides to amino acids. The aa sequences of *Pantoea* PNA 97-1R (NCBI - CP020955.2). and *Enterobacter* EcWSU1 (NCBI – CP002887.1) were compared by alignment with the “Geneious alignment” function. The percent pairwise identity was recorded.

#### Generating false color luminescence images.

Luminescence false color images were merged in ImageJ to create compound images of luminescence and brightfield captures. The TIFF file of the long exposure capture (luminescence) and brightfield capture were opened with ImageJ Fiji release. The images were merged (image→ mergechannels→ [select blue for brightfield and yellow for luminescence]). The output file was saved in PNG format for publication images.

#### Calculating 48 h thiosulfinate growth response curves.

The relative stock concentration of garlic, red onion, and allicin stock solutions in LB were used to estimate the volumes of stock solutions needed to create ten media solutions of each treatment between 0-200 µM. The actual concentration of each solution was measured using the 4-MP assay. The LB thiosulfinate dilution series solutions were added to bioscreenC plates, inoculated with PNA 97-1R WT and ∆*alt* and incubated for 48 h. The raw OD_600_ readings were normalized by subtracting initial readings from each time point. The time at which half the maximum absorbance over 48 h was recorded for that concentration (rounding up or down if the reading was not recorded). The hour response values were molar thiosulfinate concentrations were imported into Microsoft excel. A linear regression based off of the allicin supplemented LB response curve was plotted. 95% confidence intervals were added around the linear regression. Thirty data points were used for each solution to generate a linear regression with 95% confidence intervals using the ggplot2 package. The resulting regressions were plotted in excel along with the data points.

### Supplementary Tables

Table S1. Genes in the *alt* cluster of *P. ananatis* PNA 97-1R. Gene, proposed gene name; locus_tag, NCBI locus tag from version CP020945.2; HMMER Family matches, closest match with amino acid product query.

| **Gene** | **locus_tag** | **NCBI Annotation** | **HMMER Family Matches** | **E-value** | **Citations** |
| --- | --- | --- | --- | --- | --- |
| *altB* | B9Q16_23170 | SDR family oxidoreductase CDS | Tyrosine-dependent oxidoreductases | 1.10E-04 | (9) |
| *altC* | B9Q16_23165 | DsbA family oxidoreductase CDS | DsbA-like | 7.00E-02 | (10) |
| *altA* | B9Q16_23160 | alkene reductase CDS | FMN-linked oxidoreductases | 4.55E-08 | (11) |
| *altD* | B9Q16_23155 | thiol reductase thioredoxin CDS | Thioltransferase | 1.20E-02 |  |
| *altE* | B9Q16_23150 | carboxymuconolactone decarboxylase family protein CDS | CMD-like | 1.30E-02 | (12) |
| *altR* | B9Q16_23145 | TetR/AcrR family transcriptional regulator CDS | Tetracycline repressor-like, C-terminal domain | 1.50E-03 | (13) |
| *altG* | B9Q16_23140 | hypothetical protein CDS | Thioltransferase | 1.50E-03 |  |
| *altH* | B9Q16_23135 | DMT family transporter CDS | Multidrug resistance efflux transporter EmrE | 1.50E-02 | (14) |
| *altI* | B9Q16_23130 | aminotransferase class I/II-fold pyridoxal phosphate-dependent enzyme CDS | Cystathionine synthase-like | 1.21E-05 |  |
| *gorB* | B9Q16_23125 | gorA CDS | FAD/NAD-linked reductases, N-terminal and central domains | 5.93E-07 |  |
| *altJ* | B9Q16_23120 | OsmC family protein CDS | Ohr/OsmC resistance proteins | 3.10E-04 | (15, 16) |

Table S2. General cloning, deletion, and labeling strains and vectors used in this study.

| ***E. coli* Strain** | **Purpose:** | **Source:** |
| --- | --- | --- |
| MaH1 | *attTn7 pir116* R6K replicon plasmids,  DH5α derivative | (1) |
| RHO5 | *pir116,* DAP-dependent conjugation strain, SM10 derivative | (1) |
| DB3.1 | Gateway cassette maintence strain (ccdB toxin resistant) | Invitrogen |
| DH5α | General plasmid cloning strain | (17) |
| RHO3 pTNS3 | DAP dependent conjugation strain SM10 derivative with Tn*7* transposase helper plasmid (Amp^R^) | (18) |
| Plasmid Vector |  |  |
| pR6KT2 | R6K-based suicide vector used for allelic exchange, *sacB*, GmR, *gus* | (1) |
| pR6KT2G | Gateway-derivative of pR6KT2 with BP clonase compatible cassette, *sacB*, Gm^R^, *gus,* Cm^R^ | This study |
| pDONR221 | Gateway BP clonase compatible cloning vector | Invitrogen |
| pBS46 | Gateway expression vector, pBBR1-MCS5 derivative | (19) |
| pBS46::EV | Empty expression vector pBS60 | (19) |
| pCPP5242 | Source of FRT-flanked Sp^R^ resistance cassette for marked deletions | (20) |
| pTn7PA143LuxFK | miniTn*7* Lux labeling | This study |

| **Name** | **Sequence (5’ to 3’)** |
| --- | --- |
| **cloning Sequences** | |
| GtwyCst_NotF | TAATGCGGCCGCGTTAACGCTAGCATGGATGT |
| GtwyCst_NotR | TAATGCGGCCGCTCAGAGATTTTGAGACACGG  (NotI sites area underlined) |
| attB1 | GGGGACAAGTTTGTACAAAAAAGCAGGCTTA |
| attB2 | TACCCAGCTTTCTTGTACAAAGTGGTCCCC |
| soe.ar | TAACTCGAGTGCCTAGGTATGGTACCT  (XhoI, AvrII, and KpnI sites are underlined) |
| soe.bf | AGGTACCATACCTAGGCACTCGAGTTA  (KpnI, AvrII, and XhoI sites are underlined) |
| pCPP5242_SpF | TAATGCTAGCGCCAAGCTTGCATGCAGATT |
| pCPP5242_SpR | TAATGCTAGCACACAGGAACACTTAACGGCT  (NheI sites are underlined) |
| **pR6KT2G sequencing primers** | |
| pR6KT2+G_F | TATGCAGCGGAAAGTATACC |
| pR6KT2+G_R | ACAGGCTTATGTCAATTCGA |
| pR6KT2G_F | GTCTTAAGCTCGGGCCCC |
| pR6KT2G_R | GGGATATCAGCTGGATGGC |
| **OVR A knockout** | |
| OVRA.A.F | attB1-GATTCGTTGTAGCCGCTGTTTT |
| OVRA.A.R | soe.ar-ATCATCACTGTAAGGGGGAGGA |
| OVRA.B.F | soe.bf-TTTTGCCGAGTCCGAAATTGAC |
| OVRA.B.R | attB2-TTTATCTTTTCGCCGAACAGCC |
| OVRA.outF | TCGTTTTCAGTCTGAAATACAGT |
| OVRA.outR | AACCAGCTCATGACTTACCAGA |
| **OVR B knockout** | |
| OVRB.A.F | attB1-TGGTACACATCGTTTATCCGCT |
| OVRB.A.R | soe.ar- TGGTACACATCGTTTATCCGCT |
| OVRB.B.F | soe.bf- TGCGGAATTTACCTAGCGAGAG |
| OVRB.B.R | attB2-TCACCATTTCCCGCATGAAAAC |
| OVRB.outR | TCATCATTCCAGGCTCAGCG |
| OVRB.outF | GCGTTCTCTTGCCTCTGAAG |
| **OVR C knockout** | |
| OVRC.A.F | attB1- GCGTTCTGTTTGGGATGAGAAC |
| OVRC.A.R | soe.ar- ACTCTGCCTGACTTGAACCTTT |
| OVRC.B.F | soe.bf- ACTGTTATCAGGCTGATGACCG |
| OVRC.B.R | attB2- AGTGAGTGGGATTGATACCTGC |
| OVRC.outF | AAACATATAGCTGACCGTCTG |
| OVRC.outR | GGCGAACGACATCAGCTTTT |
| **OVR D knockout** | |
| OVRD.A.F | attB1- ACTGGCTACTGAGAAAAAGCGA |
| OVRD.A.R | soe.ar- ATTACCGGGCCTTTACTCTGTG |
| OVRD.B.F | soe.bf- TGGTGAAGCGCTATTTTTATGAACA |
| OVRD.B.R | attB2- CGACCTCCGAGAATGTTTTCAG |
| **alt knockout** |  |
| OVRApt1.A.F | attB1- CCGTTCCTCAGTCCCAAGAG |
| OVRApt1.A.R | soe.ar- AAAGCCTGACAGACTGCTCC |
| OVRApt1.B.F | soe.bf- ACGCTGATACGTACTGCCTG |
| OVRApt1.B.R | attB2- TGCGATGGCACTTTTCAAGC |
| OVRApt1.outF | ATATCCGACACGTTGACGCA |
| OVRApt1.outR | GAAGTAGCCGGTCCCAGATG |
| **altR knockout** | |
| altR Fusion ordered | attB1- CP020945.2_40,909-41,649- soe.bf- CP020945.2_42,272-43,123-attB2 [IDT order] |
| altR.outF | AATTAATTATACTGTTGCTT |
| altR.outR | ATTTTTCAGGGACAGTGAGC |
| **pepM knockout** | |
| pepM.A.F | attB1- GCCACCAACGTTATCTGTCC |
| pepM.A.R | soe.ar- ATAAACACCTTTGATGCGTA |
| pepM.B.F | soe.bf- AAAGTGGCGTAAATGCAGCG |
| pepM.B.R | attB2- TGAGGGATCAGTGCGTTGAG |
| pepM.outF | ACCCGGAAACATATCACGGT |
| pepM.outR | CTGGTAATGGTCCTTCAGGA |

Table S3. Primers used for allelic exchange deletion.

Primers used for constructing the gateway compatible version of pR6KT2, amplifying the FRT flanked sp cassette, and constructing deletion vectors. Gateway cloning sequences and overlap-extensions are included.

| **Name** | **Sequence (5’ to 3’)** |
| --- | --- |
| **altB-A** | |
|  | attB1- GGAGCAGTCTGTCAGGCTTT |
|  | attB2- GCCCCTGAGTCTCGACATAA |
| **altD-G** | |
|  | attB1- CGGGTTTATGGGTGGCACTA |
|  | attB2- AGAACGACAGCACCGCTTAA |
| **altH-I** | |
|  | attB1- TCCCTGCGTAAAAAGCTCGT |
|  | attB2- CATGGCTGCACGGTTAATCG |
| **altR** | |
|  | attB1- CCGAGGCGCTGCATTAATTC |
|  | attB2- CGCAGTACTGTTTCGTGCAG |
| **altB-G** | attB2-PasI-CP020945.2_38,002-42,994—PasI-attB1[Genescript order] |
| **altH-J** | attB2- CP020945.2_42,575-47,463-attB1[Genescript order] |
| **altB-J sequence confirmation primers** | |
| altFull1_F | CAGACGCTCAGAGAGACGAC |
| altFull1_R | GAAACCAGGACAGCAGAGCT |
| altFull2_F | ACCTGAAGCAGCCACTCATC |
| altFull2_R | TAATCTCAATCCGGGCGCTC |
| altFull3_F | GAGCGCCCGGATTGAGATTA |
| altFull3_R | GTCCGGACAGCCAAAGAGTT |
| altFull4_F | ACGAGCTTTTTACGCAGGGA |
| altFull4_R | CAGAAACAAACTCACCCGCG |
| altFull5_F | GCTTTCAGCTGAAACGGCAA |
| altFull5_R | GCTTCCGGCCATACCTGTTA |
| altFull6_F | GTCTGCTGATTACGGAGGGG |
| altFull6_R | ACAGACCAGATCGGACCAGA |
| altFull7_F | GCAACGCTGAGCTGATGATG |
| altFull7_R | AAGCAGGAGAATCACCCAGC |

Table S4. Complementation primers and Genescript order.

| Insert | Plasmid | Host Strain | Source |
| --- | --- | --- | --- |
| ∆OVRA | pR6KT2G | *E.coli* RHO5 | This study |
| ∆OVRB | pR6KT2G | *E.coli* RHO5 | This study |
| ∆OVRC | pR6KT2G | *E.coli* RHO5 | This study |
| ∆OVRD | pR6KT2G | *E.coli* RHO5 | This study |
| ∆alt | pR6KT2G | *E.coli* RHO5 | This study |
| ∆pepM | pR6KT2G | *E.coli* RHO5 | This study |
| ∆altR | pR6KT2G | *E.coli* RHO5 | This study |
| altB-J | pDONR221 | *E.coli RHO5* | This study |
| altB-G | PUC57 w *attB* sites | *E. coli* DH5α | Genscript (This study) |
| altB-G | pDONR221 | *E. coli* DH5α | This study |
| altH-J | PUC57 w *attB* sites | *E. coli* DH5α | Genscript (This study) |
| altH-J | pDONR221 | *E. coli* DH5α | This study |
| altB-A | pDONR221 | *E. coli* DH5α | This study |
| altD-G | pDONR221 | *E. coli* DH5α | This study |
| altH-I | pDONR221 | *E. coli* DH5α | This study |
| altB-J | pBS46 | *E. coli* DH5α | This study |
| altB-G | pBS46 | *E. coli* DH5α | This study |
| altH-J | pBS46 | *E. coli* DH5α | This study |
| altB-A | pBS46 | *E. coli* DH5α | This study |
| altD-G | pBS46 | *E. coli* DH5α | This study |
| altH-I | pBS46 | *E. coli* DH5α | This study |
| altR | pDONR221 | *E. coli* DH5α | This study |
| altR | pBS46 | *E. coli* DH5α | This study |

Table S5. Deletion vectors and complementation vectors used in this study.

| Species | Strain (Genotype) | Plasmid | Source |
| --- | --- | --- | --- |
| *Pantoea ananatis* | PNA 97-1R (WT) |  | (3) |
|  | PNA 97-1R (∆OVRA) |  | this study |
|  | PNA 97-1R (∆OVRB) |  | this study |
|  | PNA 97-1R (∆OVRC) |  | this study |
|  | PNA 97-1R (∆OVRD) |  | this study |
|  | PNA 97-1R (∆OVRA/B/C/D) |  | this study |
|  | PNA 97-1R (∆OVRB/C/D) |  | this study |
|  | PNA 97-1R (∆*alt*) |  | this study |
|  | PNA 97-1R-tn7Lux (WT) |  | this study |
|  | PNA 97-1R-tn7Lux (∆alt) |  | this study |
|  | PNA 97-1R-tn7Lux (∆pepM) |  | this study |
|  | PNA 97-1R-tn7Lux (∆alt ∆pepM) |  | this study |
|  | PNA 97-1R-tn7Lux (∆alt) | pBS46::EV | this study |
|  | PNA 97-1R-tn7Lux (∆alt) | pBS46::altB-J | this study |
|  | PNA 97-1R-tn7Lux (∆alt) | pBS46::altB-G | this study |
|  | PNA 97-1R-tn7Lux (∆alt) | pBS46::altH-J | this study |
|  | PNA 97-1R-tn7Lux (∆alt) | pBS46::altB-A | this study |
|  | PNA 97-1R-tn7Lux (∆alt) | pBS46::altB-A | this study |
|  | PNA 97-1R-tn7Lux (∆alt) | pBS46::altD-G | this study |
|  | PNA 97-1R-tn7Lux (∆alt) | pBS46::altH-I | this study |
|  | PNA 97-1R-tn7Lux (∆altR) |  | this study |
|  | PNA 97-1R-tn7Lux (∆altR) | pBS46::EV | this study |
|  | PNA 97-1R-tn7Lux (∆altR) | pBS46::altR | this study |
|  | PNA 02-18 (WT) |  | this study |
|  | PNA 02-18- tn7Lux |  | this study |
|  | PNA 02-18- tn7Lux | pBS46::EV | this study |
|  | PNA 02-18- tn7Lux | pBS46::altB-J | this study |
|  | PNA 200-3 (WT) |  | this study |
| *P. stewartii* susbp. indologens | PNA 03-3 (WT) |  | (21) |
|  | PNA 14-12 (WT) |  | (21) |
| *E.coli* | DH5α |  | (17) |
|  | DH5α | pBS46::EV | this study |
|  | DH5α | pBS46::altB-J | this study |
| *E. cloacae* | EcWSU1 |  | (22) |
|  | UW5 |  | (23) |

Table S6. *Panteoa*, *E. coli,* and *E. cloacae* strains used in this study.

### Supplementary Figures

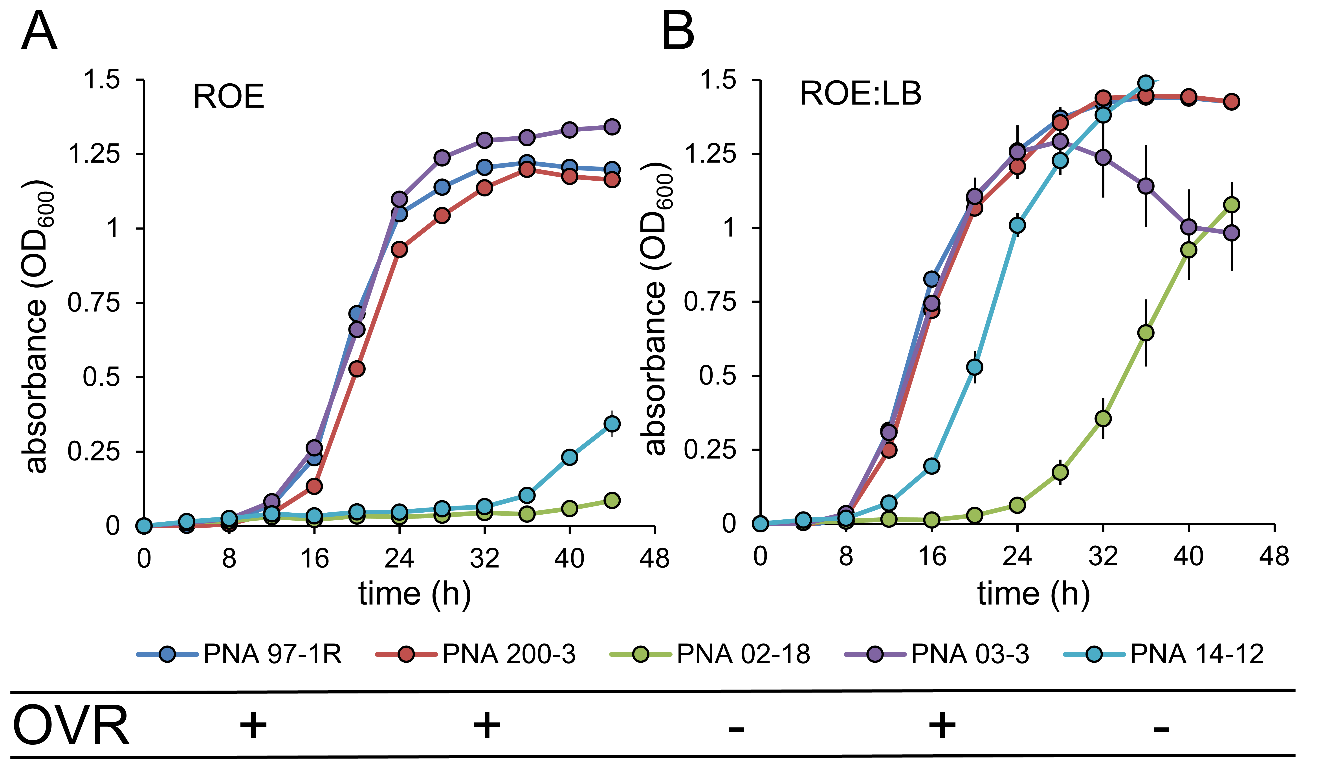

Fig S1. Growth of natural variant *Pantoea* isolates in aqueous red onion extract (ROE) and ROE amended with LB. (*A*) Growth of isolates (change in OD_600_) in ROE. (*B*) Growth of isolates in ROE:LB. This experiment was repeated three times with similar results (*N=6*). Error bars represent ±SE.

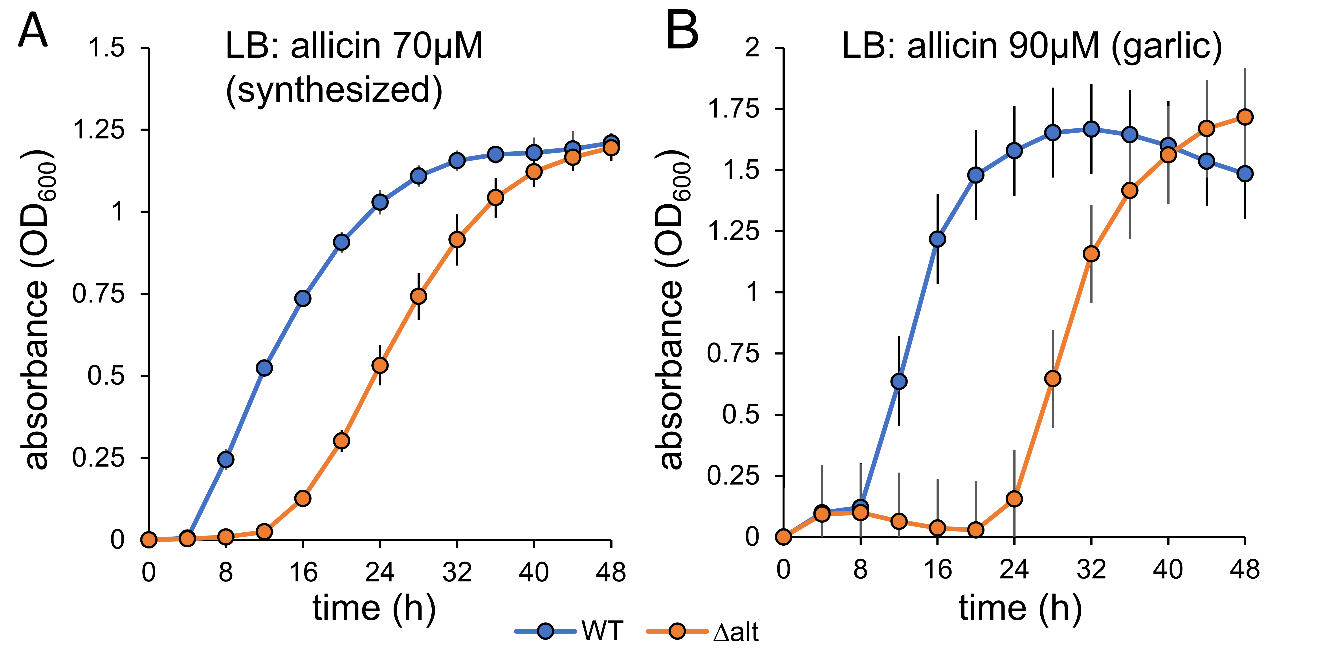

Fig S2. Growth of PNA 97-1R ∆*alt* in LB broth amended with allicin. Bioscreen growth of strains (change in OD_600_), PNA 97-1R WT and PNA 97-1R ∆alt. (*A*) synthesized allicin amended LB. (*B*) garlic extract amended LB. Allicin concentration determined with 4-MP assay. This experiment was repeated three times with similar results (N=4). Error bars represent ±SE.

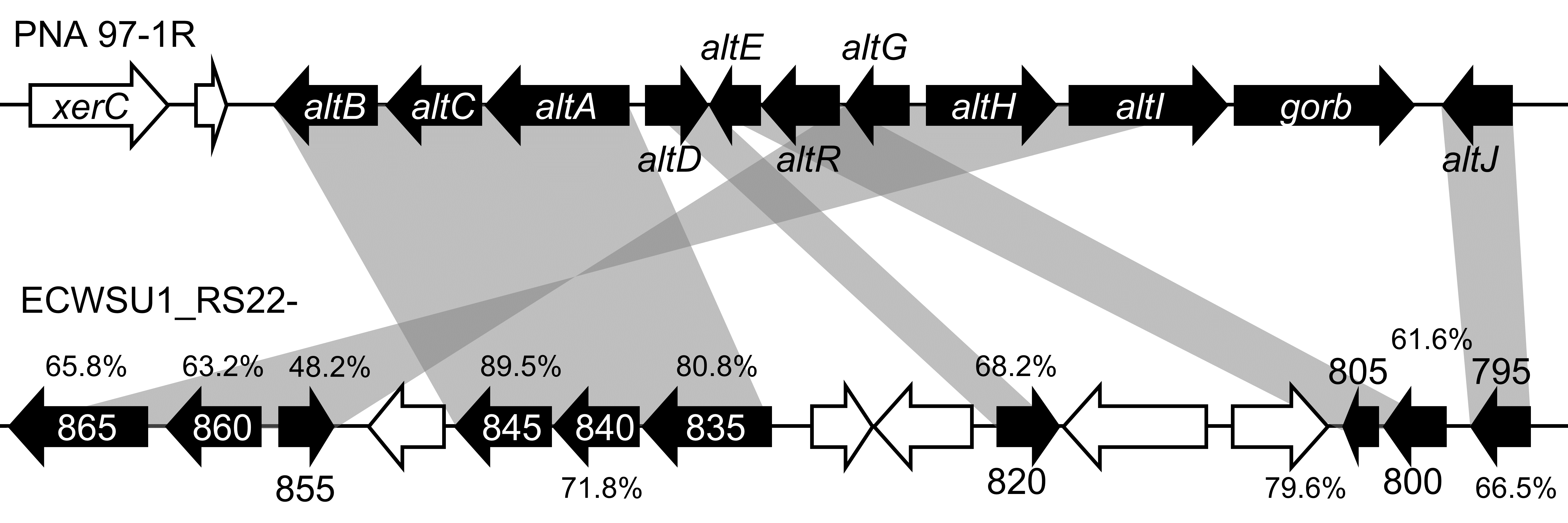

Fig S3. PNA 97-1R *alt* cluster shares homology with a plasmid-borne gene cluster of onion rot pathogen *Enterobacter cloacae* EcWSU1. Percent amino acid identity is depicted above homologous EcWSU1 genes. NCBI locus numbers are included for EcWSU1 genes.

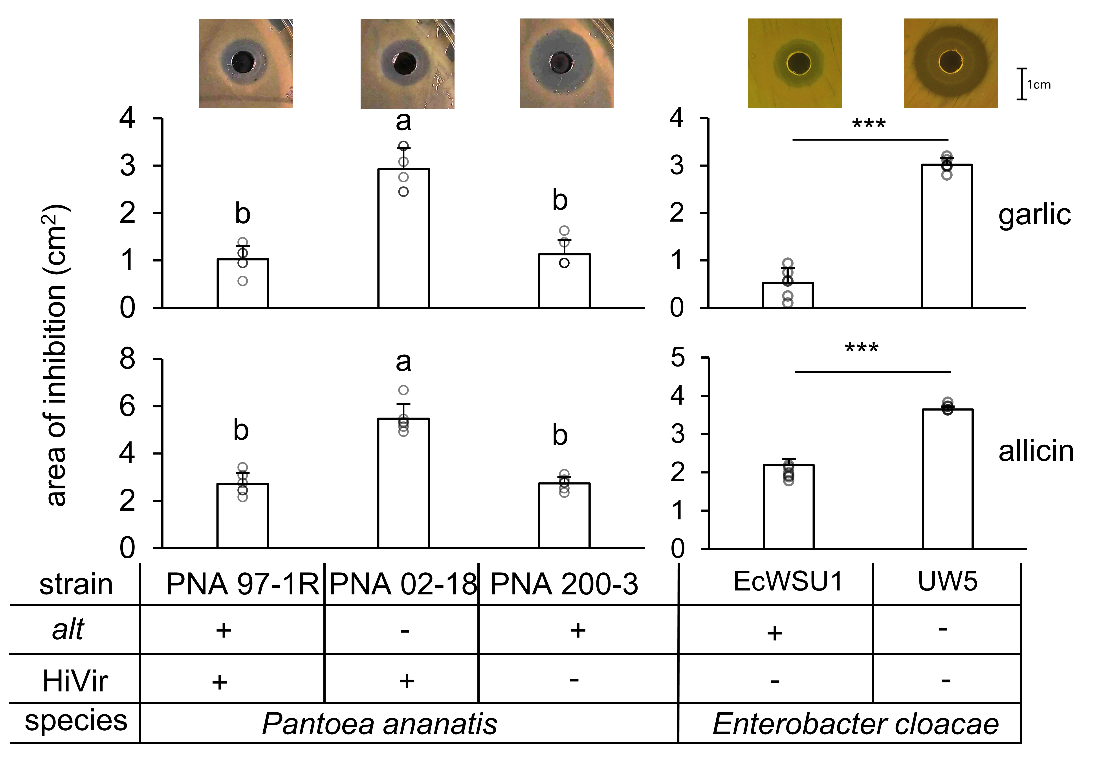

Fig S4. Natural variants of *P. ananatis* and *Enterobacter* *cloacae* with *alt*-like genes have increased tolerance to allicin and garlic extracts. Area of inhibition for garlic extract (top) and allicin (bottom) is represented. Representative zones of inhibition for allicin above corresponding bars. The data from one of three independent replicates is presented (*N=6,* one-way ANOVA followed by Tukey´s post-test) (*N=6*, t-test, ***p<0.0001). Error bars represent ±SD.

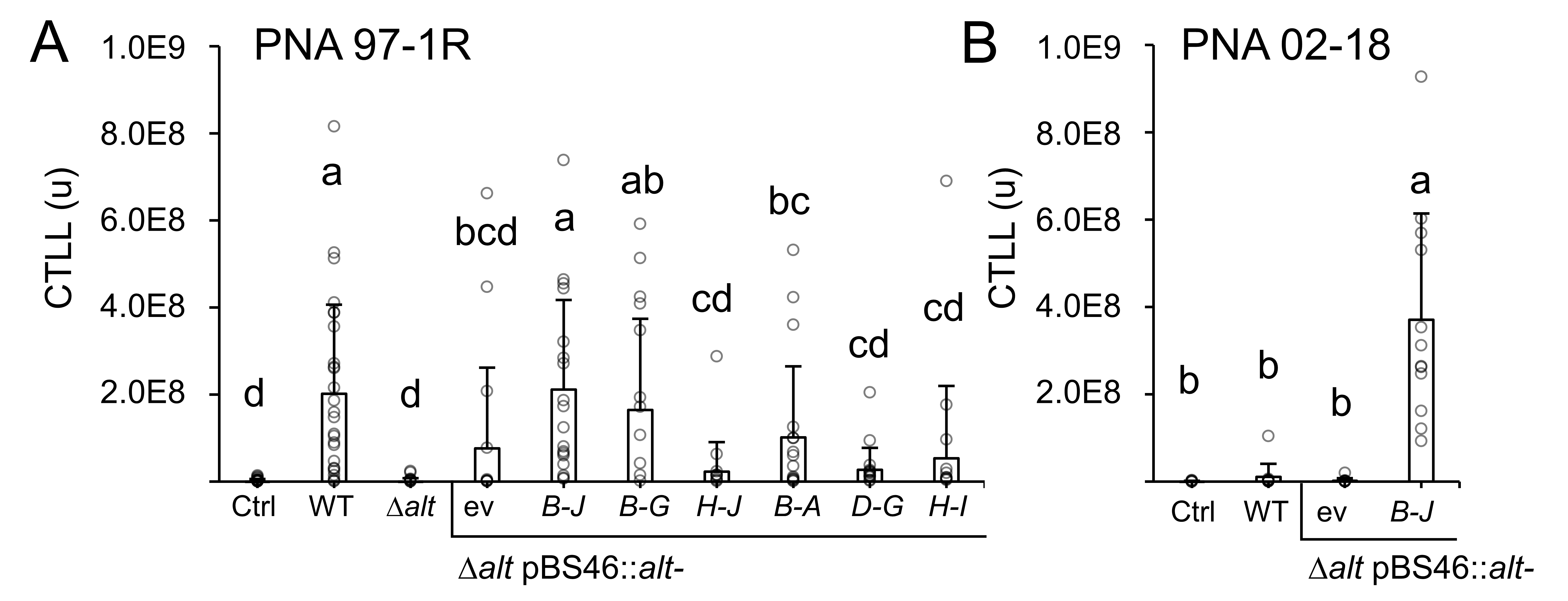

Fig S5. Calculated total lesion luminescence (CTLL) of complementation constructs in WT and heterologous *Pantoea* background. Values were quantified by defining the lesion area in stacked images and quantifying relative pixel intensity in imageJ. CTLL= integrated density – (area of lesion X mean luminescence of background readings, ctrl). (*A*) CTLL of PNA 97-1R nested complementation constructs. The data from six independent replicates is presented (*N=18,* one-way ANOVA followed by Tukey´s post-test). (*B*) CTLL of PNA 02-18 expressing *altB-J.* The data from three independent replicates is presented (*N=12,* one-way ANOVA followed by Tukey´s post-test).

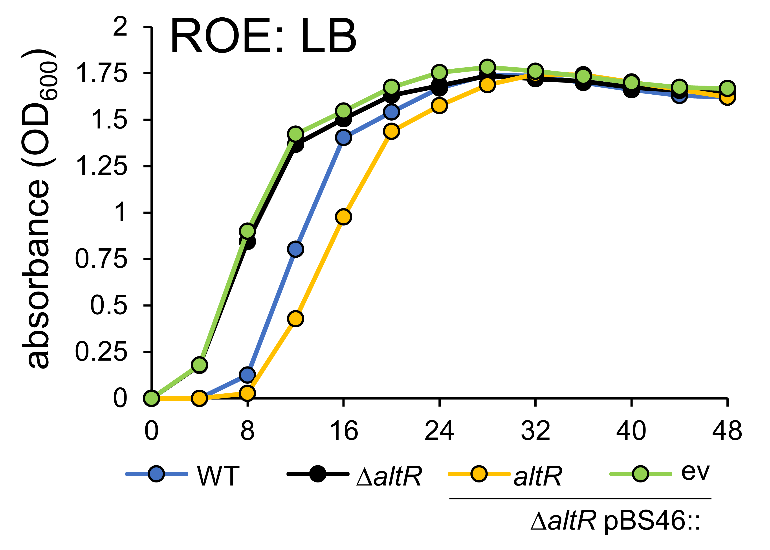

Fig S6. Growth curve of *altR* mutants in ROE:LB. Bioscreen growth of isolates (change in OD_600_), ev = empty vector. This experiment was repeated three times with similar results (*N=6*). Error bars represent ±SE.

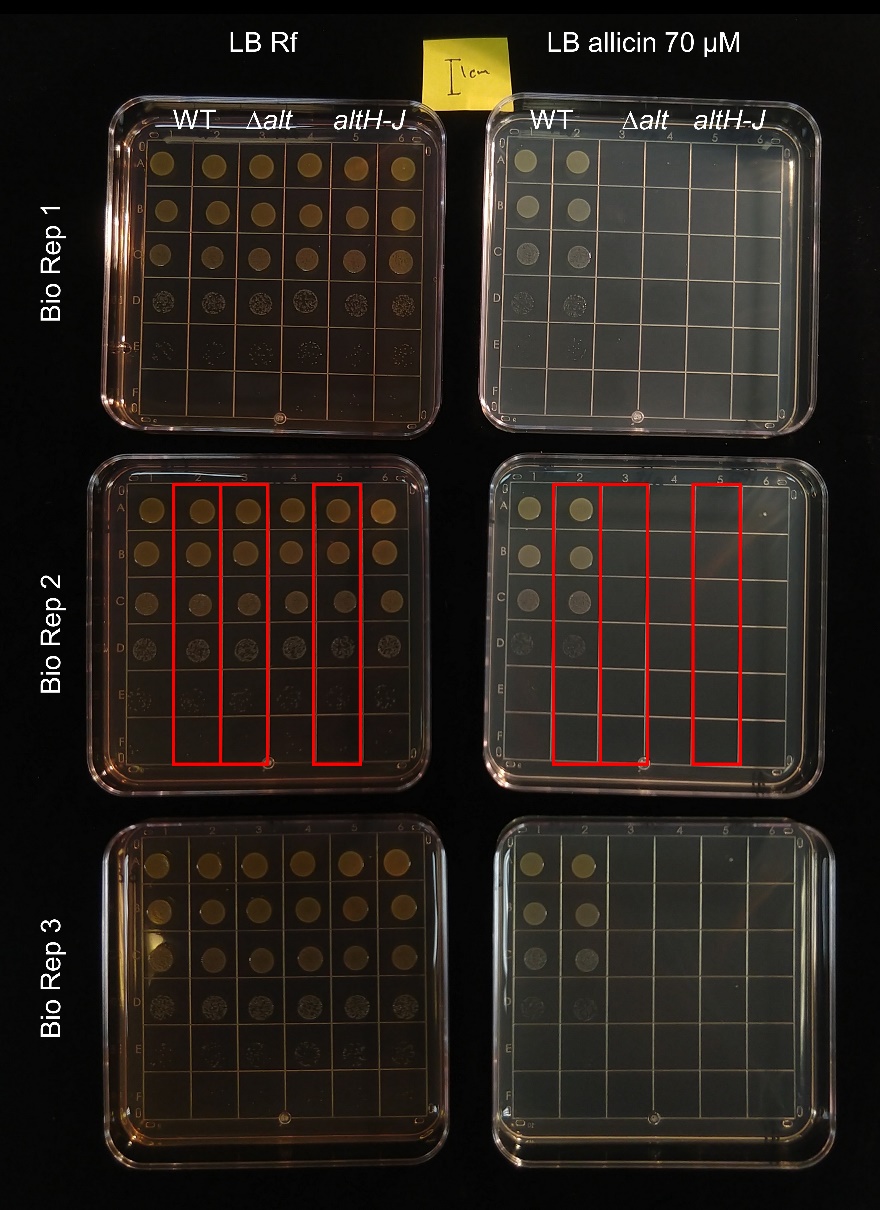

Fig. S7. WT, ∆*alt*, *altH-J* ten-fold serial dilution on LB rifampicin and LB allicin amended plates. Fig. 5 cropped panels highlighted in red.

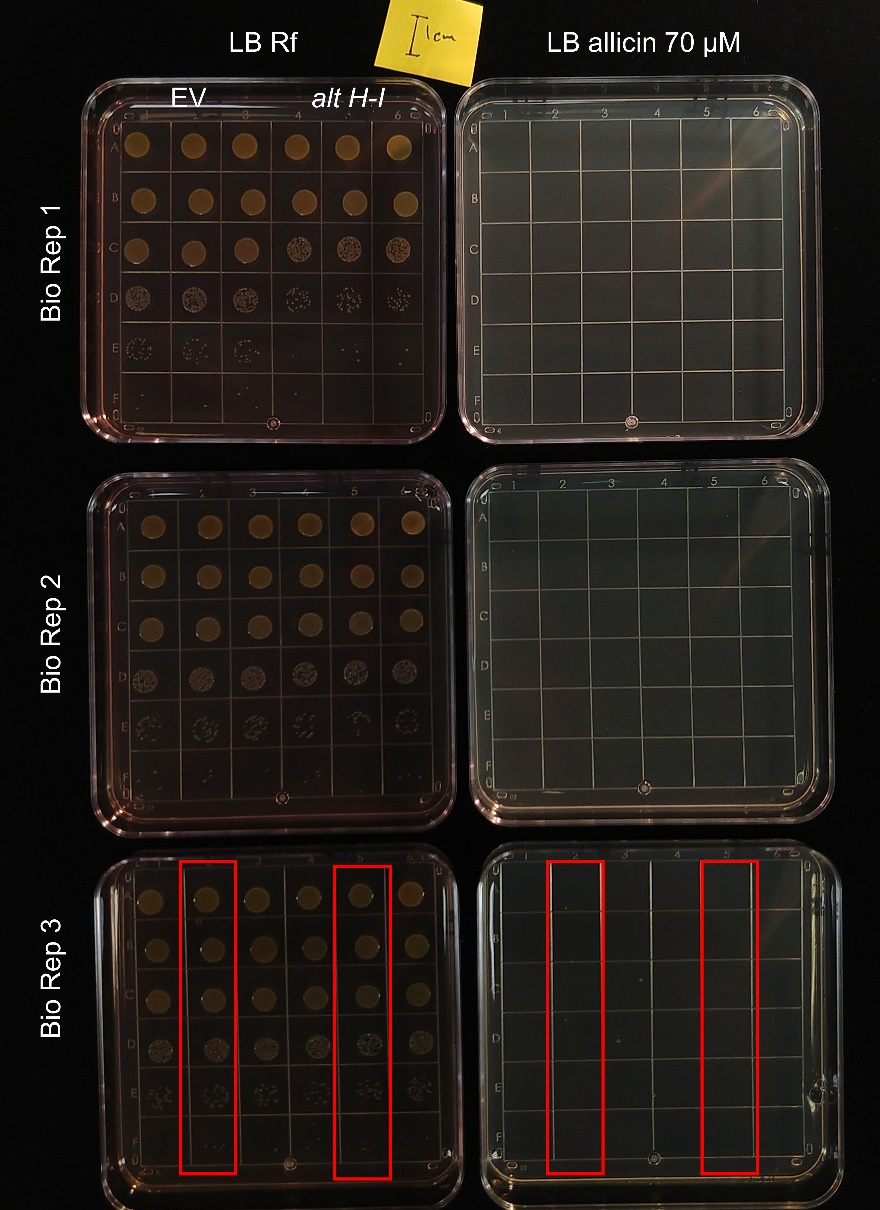

Fig S8. WT, ∆*alt*, *altH-J* ten-fold serial dilution on LB rifampicin and LB allicin amended plates. Fig. 5 cropped panels highlighted in red.

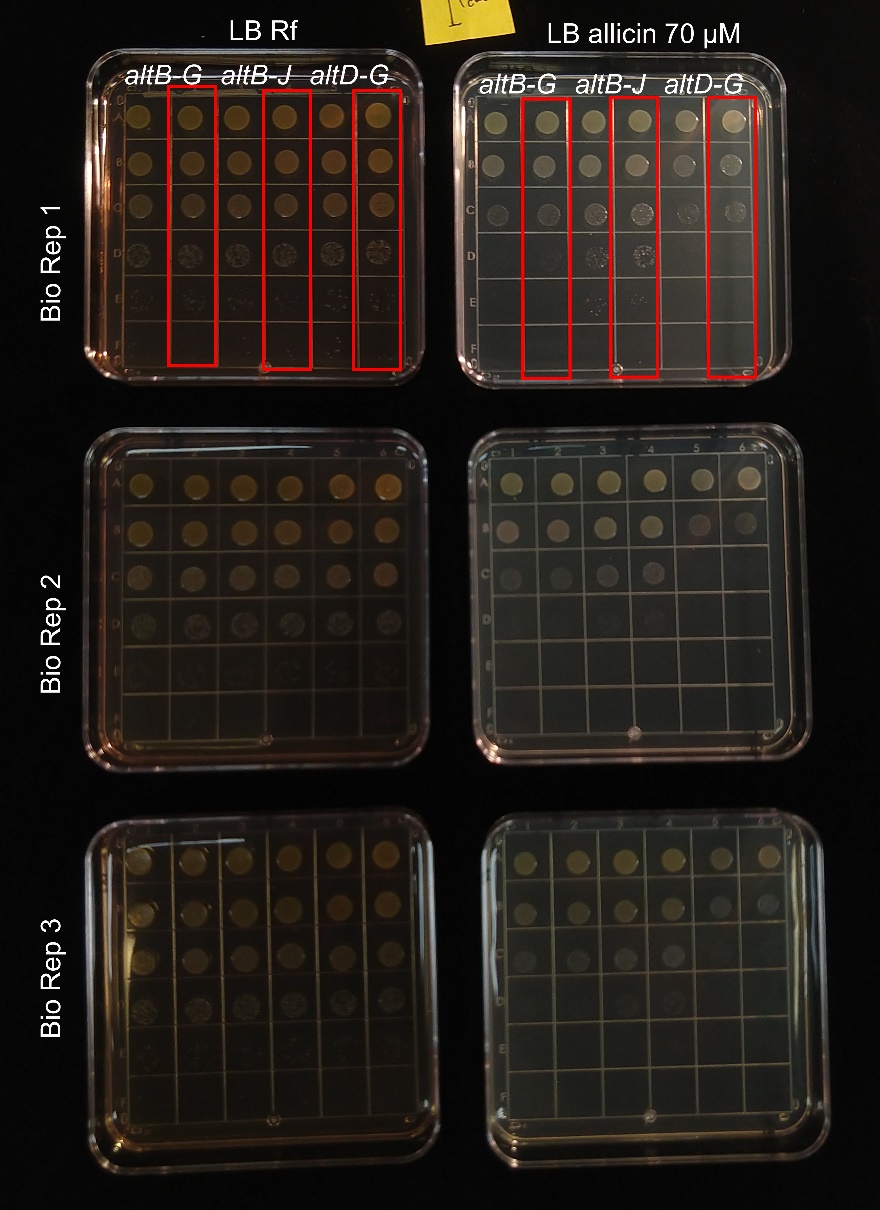

Fig S9. WT, ∆*alt*, *altH-J* ten-fold serial dilution on LB rifampicin and LB allicin amended plates. Fig. 5 cropped panels highlighted in red.

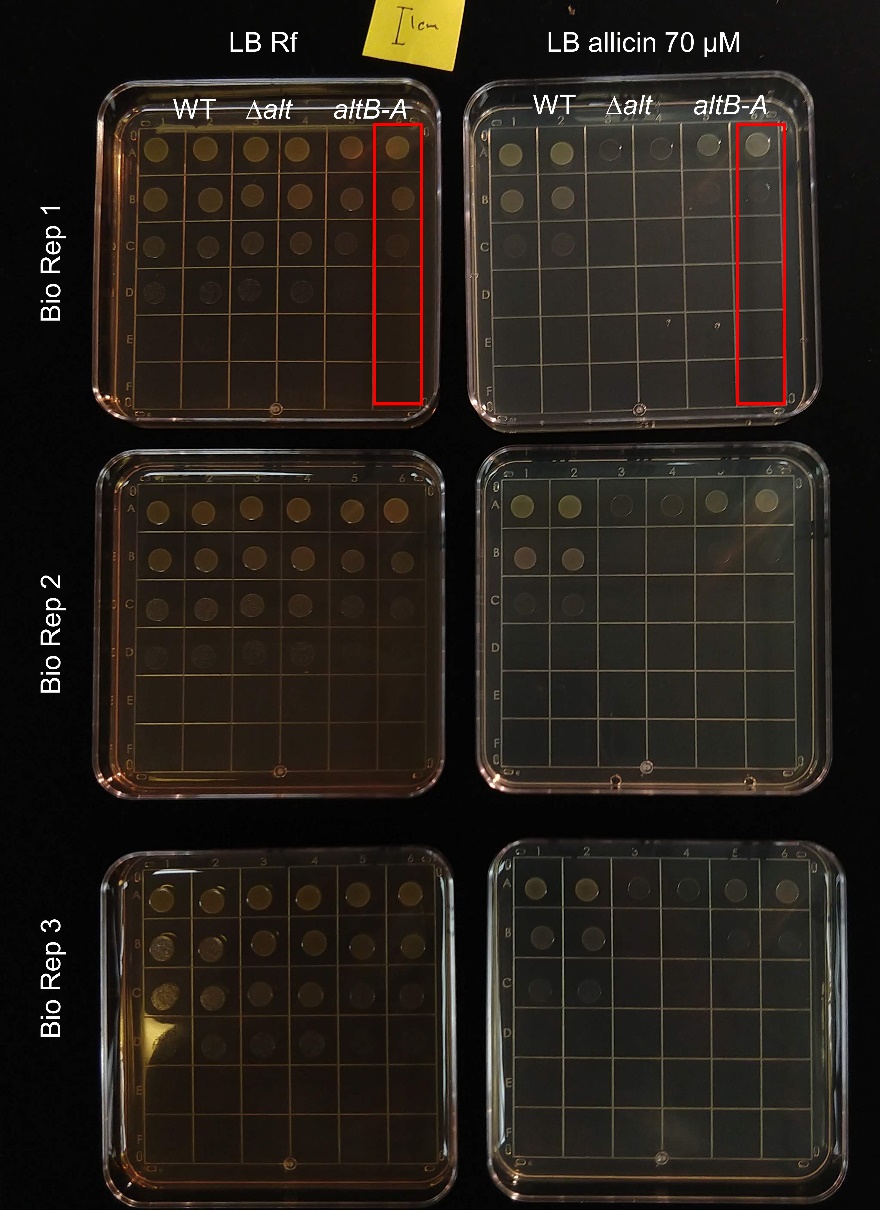

Fig S10. WT, ∆*alt*, *altH-J* ten-fold serial dilution on LB rifampicin and LB allicin amended plates. Fig. 5 cropped panels highlighted in red.

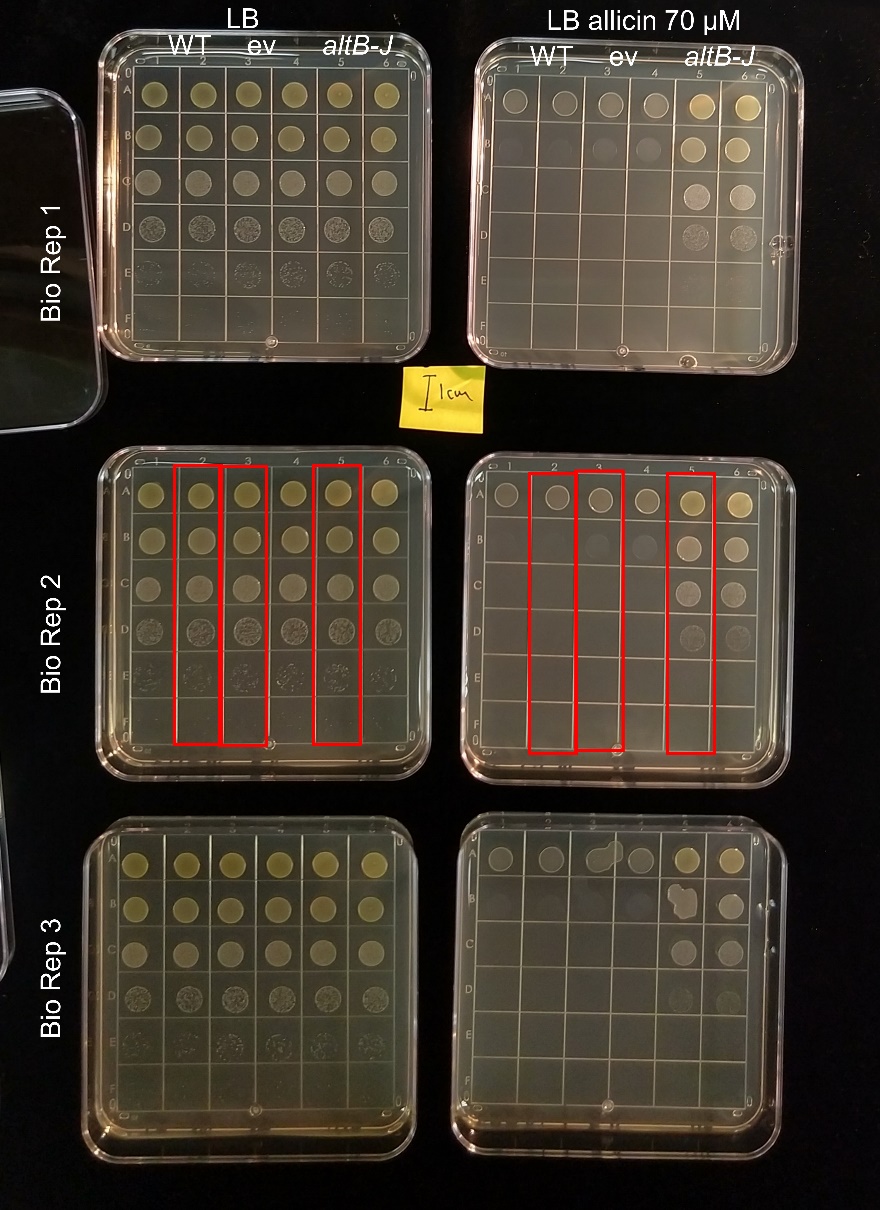

Fig S11. *P. ananatis* PNA 02-18 WT, ev, and *altB-J* ten-fold serial dilution on LB and LB allicin amended plates. Fig. 6 cropped panels highlighted in red.

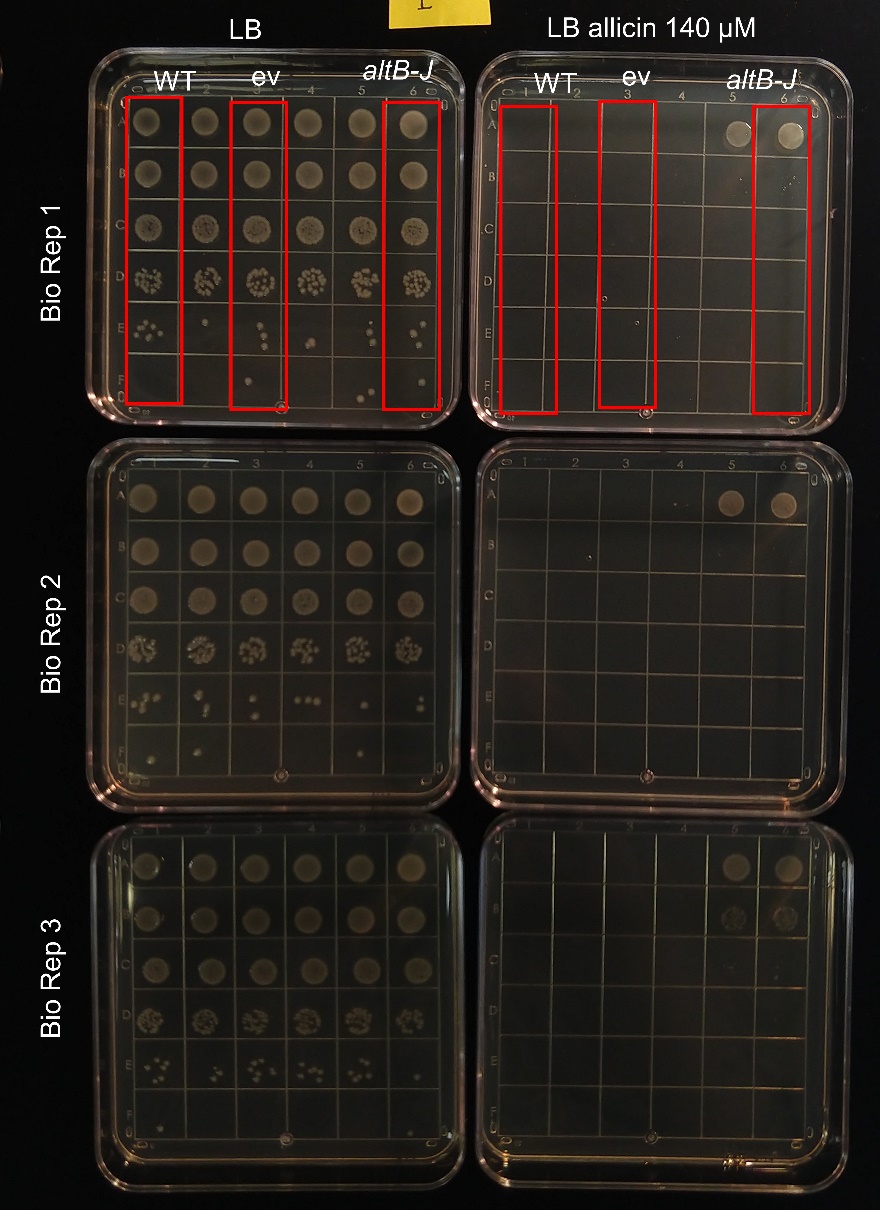

Fig S12. *E.coli* DH5α WT, ev, and *altB-J* ten-fold serial dilution on LB and LB allicin amended plates. Fig. 6 cropped panels highlighted in red.
